# NG2-targeting macrophages inhibit 3D invasion of patient-derived glioblastoma spheroids

**DOI:** 10.64898/2026.04.03.715398

**Authors:** Elena Kurudza, Sophia R.S. Varady, Daniel Greiner, James E. Marvin, Alyse Ptacek, Melanie Rodriguez, Abhinava K. Mishra, Gongping He, Gianpietro Dotti, Howard Colman, Melissa Q. Reeves, Denise J. Montell, Samuel H. Cheshier, Minna Roh-Johnson

## Abstract

Engineering macrophages with chimeric antigen receptors is emerging as a promising cancer therapeutic. Chimeric antigen receptor-expressing macrophages (CAR-Ms) engineered to recognize tumor-specific antigens have been shown to inhibit tumor growth and activate adaptive immune responses, leading to robust tumor control in animal studies. Based on this work, clinical trials have been initiated. While the trials have shown promise, challenges remain. The dynamic interactions between CAR-Ms and cancer cells and the exact mechanisms driving anti-tumor effects remain poorly defined. Defining the dynamic interactions between CAR-Ms and cancer cells will provide critical insights for optimizing future CAR-M design and improving therapeutic efficacy. We sought to directly visualize CAR-M interactions with glioblastoma cells at high-resolution and in real-time using CAR-Ms engineered to recognize Neural-Glial Antigen 2 (NG2), an antigen expressed on glioblastoma cells. Using patient-derived glioblastoma cells, we formed glioblastoma spheroids and embedded them in a 3D matrix together with CAR-Ms. Using time-lapse microscopy, as expected, we found that NG2-targeting CAR-Ms engulfed glioblastoma cells. However, excitingly, we found that NG2-targeting CAR-Ms blocked >85% of glioblastoma cell invasion in 3D. This inhibition of glioblastoma invasion was not due to a significant change in CAR-M polarization states. Together, these data suggest that NG2-targeting CAR-Ms both engulf glioblastoma cells and block glioblastoma invasive behavior.

## INTRODUCTION

Glioblastoma is the most common primary brain malignancy, making up 45.2% of malignant primary central nervous system (CNS) tumors ^1^. Despite decades of research, glioblastoma remains an incurable disease, with almost all patients succumbing within 2 years ^2^. The unique tumor microenvironment and genetic heterogeneity of glioblastoma lead to immune escape and rapid adaptation to therapy, making glioblastoma particularly difficult to treat with current approaches ^3,4^.

The tumor microenvironment, defined as a heterogeneous collection of infiltrating and resident host cells, secreted factors, and extracellular matrix, includes hallmark features such as immune cells, stromal cells, blood vessels, and neoplastic cells ^5,6^. It is well-established that the tumor immune microenvironment plays a critical role in glioblastoma development and progression ^7^. The immune system eliminates most malignant cells through immune surveillance prior to clinical recognition. When immune resistant tumor cell variants arise, tumor cells evade immune surveillance, leading to tumor development ^6,8,9^. Glioblastomas are also considered immunologically cold with an immunosuppressive microenvironment ^10–12^, limiting the durability of T cell-based immunotherapies, suggesting that alternative treatment approaches may be necessary.

Myeloid cells, including circulating macrophages and resident microglia, are key players within the glioblastoma tumor microenvironment. The myeloid compartment demonstrates extensive tumor infiltration in glioblastoma, often accounting for 50% of the tumor mass ^13,14^. Additionally, glioblastoma demonstrates a significant increase in CD14+ myeloid cells compared to the normal brain, with active recruitment of the myeloid compartment to the tumor ^13,15–18^. It has been demonstrated that high numbers of tumor-associated macrophages are associated with more aggressive tumor behavior in glioblastoma ^19–25^, and that tumor-associated macrophage infiltration is an indicator of poor prognosis ^26–29^. Alternatively, when activated by cytokines, other immune cells, or pharmacologic interventions, these tumor-associated macrophages can differentiate to a more pro-inflammatory, anti-tumor state that promotes tumor clearance ^30–37^. Thus, macrophages possess significant plasticity and exist in a spectrum of states during glioblastoma progression.

While maximal safe resection remains the cornerstone of initial treatment in glioblastoma, non-surgical treatments are necessary to address the residual microscopic and infiltrative or recurrent disease. These surgical treatments are performed in conjunction with the current standard of care, which includes radiotherapy and subsequent chemotherapy with temozolomide ^38^. With the rapid expansion of knowledge over the past decade regarding the role of the immune microenvironment in glioblastoma, immunotherapy has emerged as a promising new treatment modality. Chimeric antigen receptor (CAR) therapy is a type of immunotherapy in which immune cells are genetically engineered to specifically recognize and bind a tumor-associated antigen, leading to a cascade of signaling that leads to tumor killing by the immune cell ^39–41^. Each CAR consists of an extracellular antigen-recognition moiety linked to an intracellular signaling domain via a hinge domain and transmembrane domain. Classically, scientists and clinicians have used T cells (CAR-T cell therapy) for immunotherapy approaches due to their innate robust ability to kill tumor cells ^42^. In glioblastoma, multiple ongoing and completed phase 1 clinical trials have explored the safety, feasibility, and efficacy of CAR-T cell therapy ^11,40,43–49^. The most common molecular targets include EGFR, HER2, and IL13Rα2.

Additionally, multi-specific CAR-T cells, targeting multiple antigens rather than just one, or successive treatments against distinct tumor antigens, are currently being tested to address intratumoral heterogeneity observed in glioblastoma ^50^. Overall, these trials have demonstrated that autologous CAR-T cells can be safely administered via peripheral and locoregional routes ^44,47^. However, CAR-T cells have demonstrated only modest efficacy in these clinical trials due to antigen modulation, immune exhaustion, and poor solid tumor infiltration ^11,40,43–49^.

Given the ability of macrophages to infiltrate solid tumors, reprogram the tumor microenvironment, and participate in antigen presentation to promote an adaptive anti-tumor response, genetically engineering a patient’s circulating macrophages into CAR-Macrophages (CAR-Ms) is a promising alternative to CAR-T cell therapy. Previous work in glioblastoma models has shown that combining CD133-targeting CAR-Ms with anti-CD47 approaches, thereby blocking the “don’t eat me” signal on cancer cells ^30,51–53^, led to significantly reduced recurrence, prolonged survival, and the induction of long-term immunologic memory in syngeneic murine models ^54^. Other studies generating glioblastoma-targeting CAR-Ms from stem cells ^55,56^ or microbe-guided CAR-Ms ^57^ have shown enhanced cytotoxicity against immortalized glioblastoma cell lines ^55,57^ and reprogramming of tumor-associated macrophages into an anti-tumor-like fate ^56,57^. CAR-Ms have not yet been tested in glioblastoma clinical trials; however, HER2-targeting CAR-Ms were recently explored in early phase 1 clinical trials in gastric and breast cancer ^58^, as well as Mesothelin-targeting CAR-Ms in ovarian, pancreatic, adenocarcinoma, and mesothelioma ^59^. These phase I clinical trials were promising in that CAR-Ms were safely administered to patients with little adverse challenges, although the treatments failed to control disease progression. The results of these clinical trials highlight the safety and tolerability of CAR-Ms for solid tumor treatment, but also the need to better understand how CAR-Ms dynamically interact with the tumor during tumor progression to further inform future design and develop more efficacious treatments.

We developed a CAR-M designed to recognize and bind Neuron Glial antigen 2 (NG2) to dissect CAR-M and glioblastoma interactions at the cellular level. NG2 is a transmembrane proteoglycan known to promote proliferation, migration, and survival of glioblastoma cells ^60–62^. Whereas prior CAR-M studies in glioblastoma have primarily established feasibility, immune activation, and tumor regression, we used NG2-targeting CAR-Ms to determine real-time, target-dependent mechanisms by which CAR-Ms interact with and restrain glioblastoma cells in 3D – demonstrating that CAR-Ms can suppress tumor growth and invasion through mechanisms that extend beyond direct phagocytosis. Specifically, using both immortalized human glioblastoma cell lines and patient-derived glioma stem cells, we demonstrate that NG2-targeting CAR-Ms recognize and phagocytose glioblastoma cells and limit tumor growth in 3D tumor spheroid models. Additionally, we demonstrate that NG2-targeting CAR-Ms robustly inhibit glioblastoma 3D invasion. These data highlight unexpected mechanisms by which CAR-Ms exert an anti-tumor effect in glioblastoma, informing further therapeutic design strategies.

## RESULTS

Given the potential of CAR-Ms for treating solid tumors, we sought to determine how CAR-Ms interact with GBM cells in real time within a 3D environment. We first identified endogenous glioblastoma cell targets that were specifically and highly expressed on the surface of glioblastoma cells, and targets in which single-chain variable fragments (scFvs) have already been identified and are currently being used in CAR-T cell trials for other cancers. We identified a candidate protein that met these criteria through analysis of publicly available transcriptomic and proteomic data ^63–65^: NG2 (Neural Glial Antigen 2), also known as CSPG4 (Chondroitin Sulfate Proteoglycan 4) and HMW-MAA (High Molecular Weight Melanoma-Associated Antigen). NG2 is highly expressed in 67% of GBM specimens ^66^ and plays a significant role in glioma formation and progression, contributing to cell migration, proliferation, and survival. The level of NG2 expression also correlates with worse GBM prognosis ^67^. Strategies inhibiting NG2 function, either through anti-NG2 antibodies ^68^ or by knocking down NG2 expression ^69^, inhibit glioblastoma growth in vivo. Anti-NG2 CAR-T cells also exhibit strong reactivity to glioblastoma cells with minimal off-target effects ^66^. Furthermore, a CAR-T cell therapy targeting NG2 is currently being used in phase I clinical trials for head and neck cancer (NCT06096038), suggesting specificity with minimal off-target effects. It remains to be determined whether NG2 is the ideal clinical target for immunotherapy approaches; however, given the high and specific NG2 expression in GBM, we sought to use this tumor antigen as a model to visualize how CAR-Ms interact with GBM cells in a 3D environment to better understand how CAR-Ms can be engineered to inhibit GBM growth and invasion.

We generated NG2-targeting CAR-Ms using primary human macrophages isolated from healthy blood donors (**Figure 1A**). Following the design principles of previous CARs ^42,70–72^, we used the NG2-targeting single-chain variable fragment (scFv) that is currently being used in CAR-T cell clinical trials for head and neck squamous cell carcinoma (NG2^763.74^-scFv). We fused the NG2^763.74^-scFv to a CD8 transmembrane sequence, and the FcR gamma intracellular signaling domain to promote downstream phagocytic responses (**Figure 1B**). Importantly, while NG2 is expressed by some healthy cell types, the antigen recognized by the specific NG2^763.74^-scFv of the chimeric antigen receptor is not detected on other cell types in the brain ^61,66,73,74^. We also included a self-cleaving P2A-EGFP-CAAX domain to mark CAR-Ms for downstream analysis (**Figure 1B**). EGFP-CAAX alone was used as a negative control. Using lentiviral transduction protocols previously established in our lab ^70,75^, we consistently achieved high rates of transduction of primary human macrophages for both NG2^763.74^-targeting CAR-Ms (herein referred to as NG2-CAR-M) and the EGFP control macrophages (**Figure 1C**).

**Figure 1:**
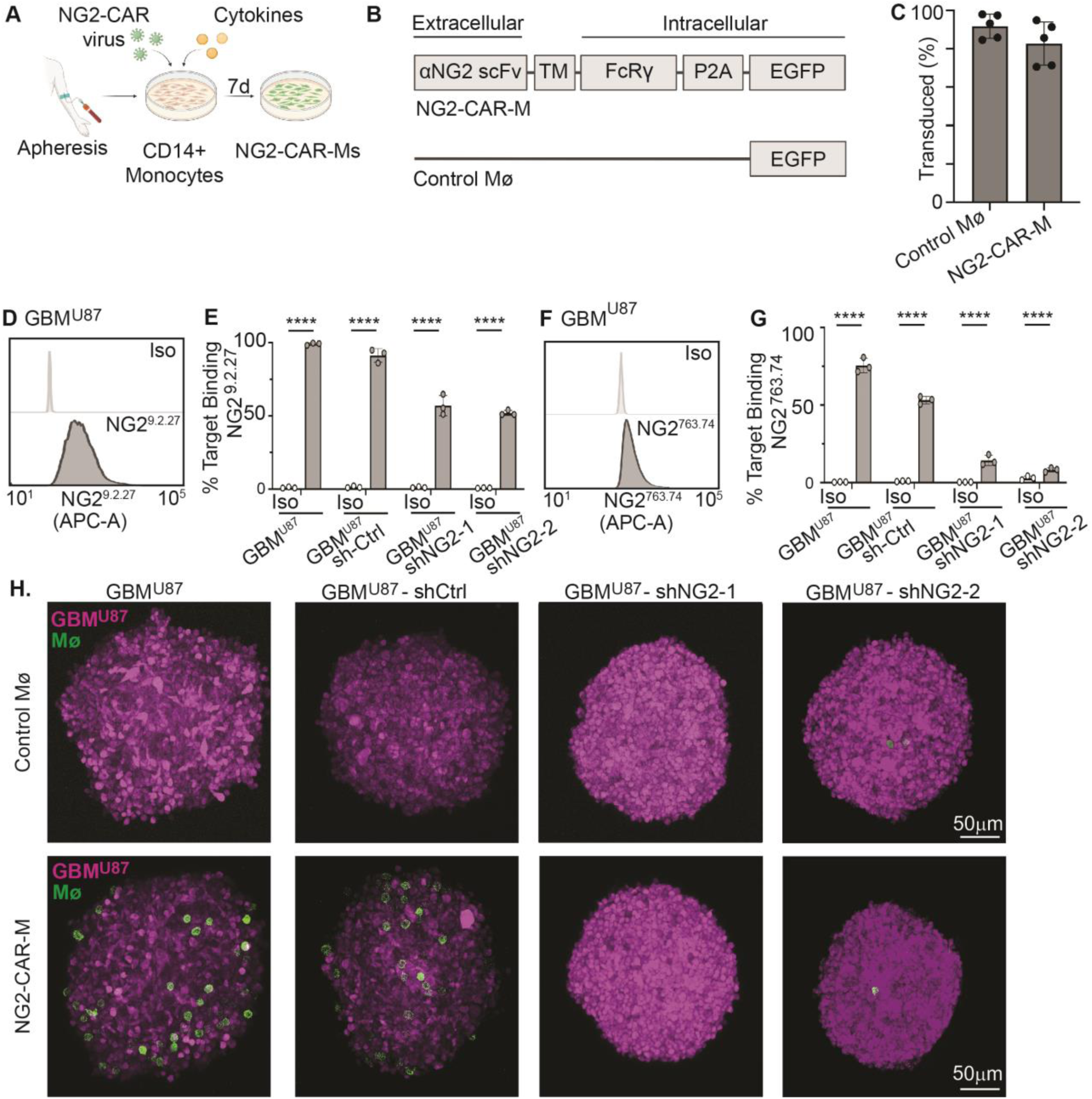
NG2-targeting CAR-Ms recognize NG2-expressing GBM cells. A. Schematic of CAR-M generation from anonymous blood donors. B. Schematic of the NG2-targeting chimeric antigen receptor constructs (top) and the EGFP control (bottom). scFv – single chain variable fragment; TM – transmembrane domain; FcRγ – FcR gamma intracellular signaling domain; P2A – Porcine teschovirus-1 2A self-cleaving peptide; EGFP – enhanced green fluorescent protein. C. Quantification of the percent of primary human macrophages that are EGFP-positive after transduction with EGFP control (Control MΦ) or the NG2-targeting CAR construct (NG2-CAR-M). D. Representative flow cytometry histogram of GBM^U87^ cells surface-stained with the NG2^9.2.27^ antibody versus an isotype control (Iso). (E) Quantification of the percent of GBM^U87^ cells positive for NG2^9.2.27^ staining versus isotype controls (Iso) when GBM^U87^ cells are untreated or treated with NG2 shRNAs (shNG2-1 and shNG2-2) versus a control shRNA (sh-Ctrl). N=3 technical replicates. 2-way ANOVA with Tukey’s multiple comparison test. **** = p<0.00001. F. Representative flow cytometry histogram of GBM^U87^ cells surface-stained with the NG2^763.74^ antibody versus an isotype control (Iso). G. Quantification of the percent of GBM^U87^ cells positive for NG2^763.74^ staining versus isotype controls (Iso) when GBM^U87^ cells are untreated or treated with NG2 shRNAs (shNG2-1 and shNG2-2) versus a control shRNA (shCtrl). N=3 technical replicates. 2-way ANOVA with Sidak’s multiple comparison test. **** = p<0.00001. H. Representative images of GBM^U87^ spheroids (magenta) cultured with control MΦ (top panels, green) or NG2-CAR-Ms (bottom panels, green). GBM^U87^ cells used to make spheroids were either untreated or treated with NG2 shRNAs (shNG2-1 and shNG2-2) versus a control shRNA (shCtrl).

We next determined whether NG2-CAR-Ms recognize NG2-expressing GBM cells. We initiated studies with a well-established immortalized human glioblastoma cell line, U87-MG, and obtained a U87-MG line that expresses RFP/firefly luciferase (Cellomics Technology), herein referred to as GBM^U87^. We first tested whether GBM^U87^ cells express NG2 on the surface of the cells by using two different NG2 antibodies: one that recognizes NG2 in an un-specified region, NG2^9.2.27^ (**Figure 1D, E**), and the other antibody that recognizes the same epitope as the NG2^763.74^-scFv used in the CAR (**Figure 1F, G**), and then compared these antibodies to an isotype control (Iso). Using flow cytometry, we found that 99.0 +/- 0.8% of GBM^U87^ cells express NG2 on the surface as detected by the NG2^9.2.27^ antibody (**Figure 1D, E**), with 75.5 +/- 4.7% of GBM^U87^ cells expressing surface NG2 that is specifically recognized by the NG2^763.74^-scFv (**Figure 1F, G**). To test for target binding specificity of the antibodies, we generated NG2 knockdown lines using shRNA approaches. We found that NG2 surface binding was significantly reduced in GBM^U87^ cells that express NG2 shRNAs (shNG2-1 or shNG2-2) compared to the non-targeting shRNA control (sh-Ctrl) (**Figure 1D-G**). These results suggest that the NG2^763.74^-scFv used in the NG2-targeting CAR recognizes NG2 expressed on the surface of GBM^U87^ cells.

We next tested whether macrophages expressing the NG2-CAR recognize and bind to NG2-expressing GBM^U87^ cells. Using the GBM^U87^ RFP-luciferase line, GBM^U87^ spheroids were made in ultra-low-attachment U-well plates over 3 days. NG2-CAR-Ms or control EGFP macrophages were added to the spheroids for 8 hours, after which the spheroids were washed, fixed, and imaged. We found that NG2-CAR-Ms adhered to the surface of GBM^U87^ spheroids within the 8-hour window. In contrast, control macrophages were not detected on the surface of GBM^U87^ spheroids, suggesting reduced or no adherence to the spheroids (**Figure 1H**, comparing “GBM^U87^” conditions in the first column). Furthermore, GBM^U87^ cells expressing NG2 shRNAs exhibited reduced numbers of NG2-CAR-Ms on the surface of the spheroid compared to non-targeting shRNA control GBM^U87^ spheroids (**Figure 1H**, shNG2-1/shNG2-2 compared to sh-Ctrl). Together, these results suggest that NG2-CAR-Ms specifically recognize and bind to NG2-expressing GBM cells.

Due to the phagocytic nature of macrophages and the use of the FcR gamma intracellular signaling domain in the CAR to induce phagocytosis, we next asked whether NG2-CAR-Ms engulf NG2-expressing GBM^U87^ cells. NG2-CAR-Ms or control macrophages were added to GBM^U87^ spheroids (**Figure 2A**). After 24 hours, we first confirmed that EGFP+ CAR-Ms successfully internalize GBM^U87^ fragments compared to control macrophages by performing Y-Z analysis of CAR-Ms that engulfed RFP+ GBM^U87^ fragments (**Figure 2B**). We then mechanically dissociated 3D co-cultures into single-cell suspensions and analyzed them by flow cytometry to quantify engulfment events by NG2-CAR-Ms compared to control macrophages (defined as CD45+, EGFP+ macrophages containing RFP+ GBM^U87^ fragments) (**Figure 2C; Figure S1**). We observed a significant amount of variability in the ability of CAR-Ms to engulf GBM^U87^ fragments (**Figure S2A**), which we attribute to PBMC donor variability since we also observed significant variability with PBMC donors in prior work ^70,76^. However, despite this variability, we still observe a significantly higher percentage of NG2-CAR-Ms engulfed GBM^U87^ fragments compared to control macrophages (**Figure S2A**). When we take into account the PBMC donor variability by normalizing engulfment rates to control for each PBMC donor, we consistently observe a significantly higher percentage of NG2-CAR-Ms that engulfed GBM^U87^ fragments compared to control macrophages (**Figure 2D**). These results suggest that NG2-CAR-Ms exhibit GBM engulfment.

**Figure 2:**
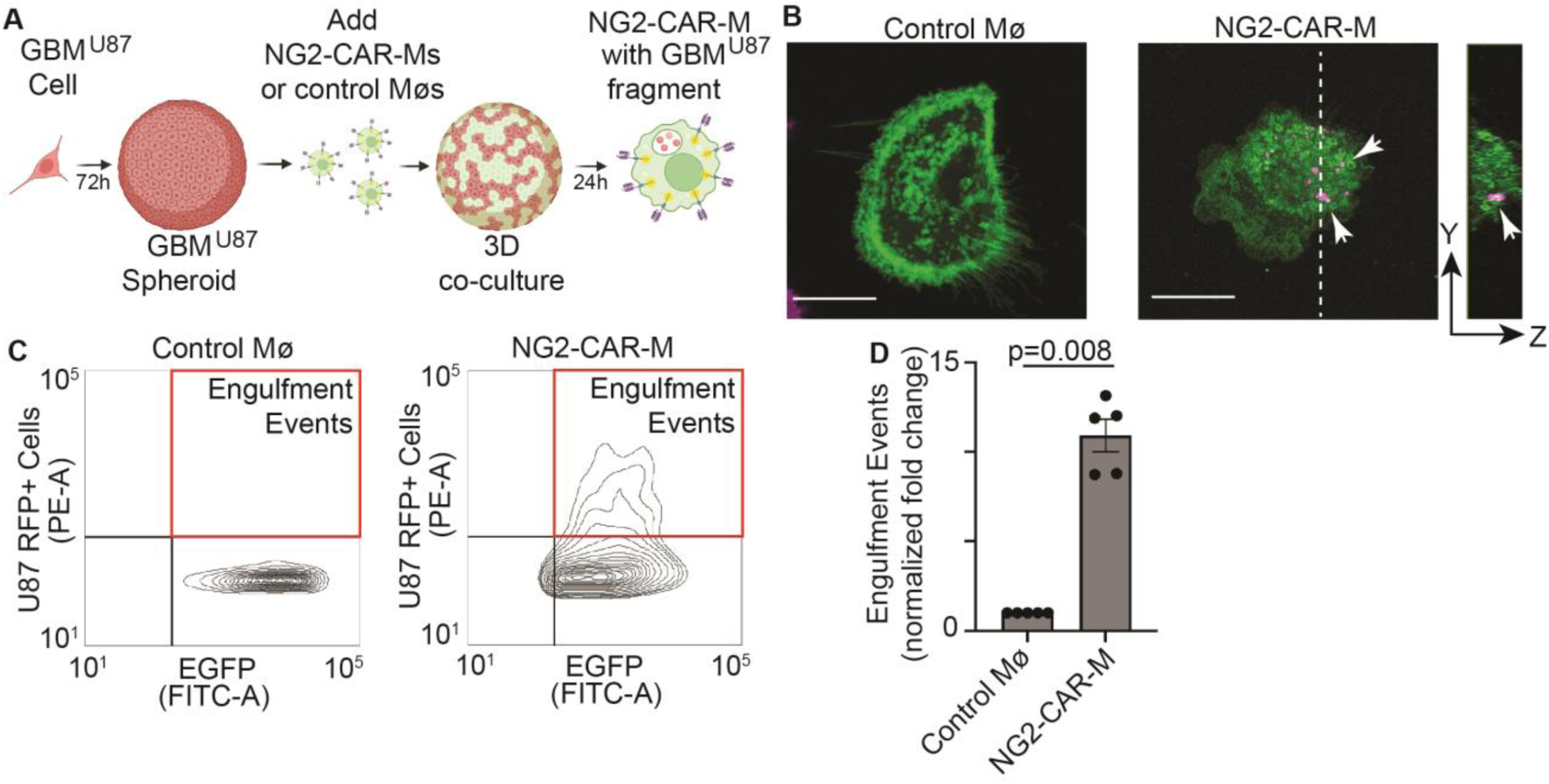
NG2-targeting CAR-Ms engulf NG2-expressing GBM cells. A. Schematic of CAR-M-mediated GBM spheroid engulfment assays. B. Representative images of Control MΦ (left) that does not contain RFP+ GBM^U87^ fragments. Representative image of NG2-CAR-M (right) that engulfed RFP+ GBM^U87^ fragments (arrows). A dotted line represents a Y-Z slice through the z-stack revealing internalization of the RFP+ GBM^U87^ fragment to the far right (arrow). Scale bar is 20 μm. C. Representative flow cytometry plot for CAR-M-mediated engulfment of GBM^U87^ cells. EGFP+ control macrophages or NG2-CAR-Ms with RFP+ GBM^U87^ fragments are quantified (red box). D. Quantification of CAR-M-mediated engulfment events from C. n=4 independent PBMC donors (biological replicates) each as a shade of gray, each with 3 technical replicates. The average of technical replicates for each PBMC donor is graphed and normalized to control macrophages, mean +/- SEM, Wilcoxon test.

The increase in GBM^U87^ engulfment by NG2-CAR-Ms versus control CAR-Ms (**Figure 2D**) suggested that NG2-CAR-Ms might inhibit, or potentially even shrink, the size of GBM spheroids. To test this hypothesis, GBM^U87^ spheroids were cultured in suspension alone (GBM alone), or co-cultured in suspension with either control macrophages or NG2-CAR-Ms, and spheroid growth was monitored over 7 days (**Figure 3A,B; Figure S2B)**. At day 7, we assessed spheroid size by quantifying the fold change in spheroid fluorescence intensity at the last time point relative to the first time point. We found that GBM^U87^ spheroids cultured with control macrophages exhibited a ∼25% decrease in spheroid size compared to GBM^U87^ spheroids cultured alone (**Figure 3C; Figure S2B**). Significantly, GBM^U87^ spheroids cultured with NG2-CAR-Ms exhibited a ∼75% decrease in size compared to GBM^U87^ spheroids cultured alone (**Figure 3C; Figure S2B**). These results suggest that NG2-CAR-Ms inhibit GBM spheroid growth. To determine whether the inhibited GBM^U87^ spheroid growth was due to changes in GBM^U87^ proliferation, we used the Ki67 proliferation marker and evaluated DNA content (DAPI). We found no difference in the percentage of GBM^U87^ cells in the G2 and Mitotic (M) phases of the cell cycle when GBM^U87^ spheroids were cultured alone or with control macrophages or NG2-CAR-Ms for 24 hours (**Figure 3D,E; Figure S3A,B)**. These results suggest that NG2-CAR-Ms do not regulate GBM proliferative capacity, and suggest that the reduced GBM spheroid growth is likely due to other mechanisms, such as increased GBM engulfment.

**Figure 3:**
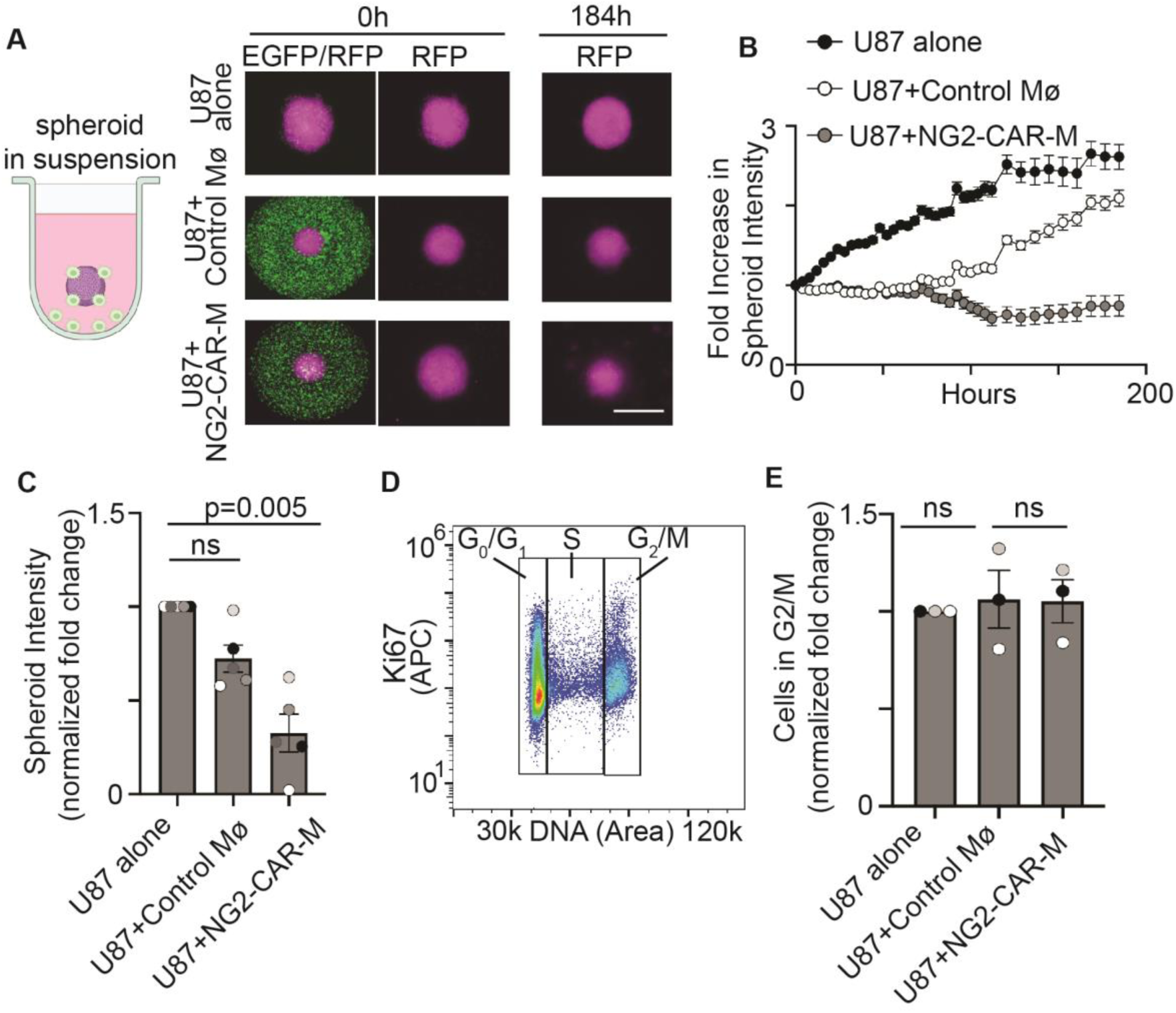
NG2-targeting CAR-Ms inhibit GBM spheroid growth. A. Schematic to the left showing spheroids in suspension. RFP+ GBM^U87^ spheroids were formed for 3 days and then EGPF+ control macrophages or NG2-CAR-M were added and spheroid growth was quantified for more 7 days. Representative images of the EGFP CAR-M channel at the first time point of addition (0h, left) and then the RFP GBM^U87^ spheroid channel showing spheroids at the first (0h, middle) and last (184h, right) time point of spheroid growth when cultured alone, or with control macrophages or NG2-CAR-Ms. Scale bar = 1mm. (B) Representative quantification of GBM^U87^ spheroid growth over time when cultured alone, with control macrophages or NG2-CAR-Ms. Data are presented as fold changes of U87 RFP fluorescence intensity at each time point relative to the initial time point. C. Quantification of data in (B) at 184 hours across conditions (n=5 independent PBMC donors (biological replicates), each with 6 technical replicates for every condition). Data are shown as fold increase in spheroid fluorescence intensity, normalized to the U87 alone control. The average of technical replicates for each PBMC donor is graphed, mean +/- SEM, Friedman’s test with Dunn’s multiple comparison. D. Representative flow cytometry analysis for Ki67 and DAPI (DNA) to categorize GBM^U87^ cells in cell cycle phases. E. Quantification of the fold change of GBM^U87^ cells in the G2/M phase of the cell cycle when cultured with control macrophages or NG2-targeting CAR-Ms, normalized to the U87 alone condition. n=3 independent PBMC donors (biological replicates). The average of technical replicates for each PBMC donor is graphed, mean +/- SEM, Friedman’s test with Dunn’s multiple comparison.

We next asked whether increasing the engulfment capacity of NG2-CAR-Ms could further inhibit the growth of GBM spheroids. We re-engineered the CAR by including a hyperactivating variant in the Rac2 GTPase, Rac2^E62K^ ^77^. The Rac2^E62K^ variant had previously been shown to increase engulfment of target cells by CAR-Ms ^78^. We created an NG2-CAR with the Rac2^E62K^ variant (NG2-Rac2^E62K^-CAR-M) and a series of control constructs (**Figure 4A**). A non-targeting Rac2^E62K^ control was created to test for the effects of the hyperactivating Rac2^E62K^ variant independent of NG2 binding (NT-Rac^E62K^-CAR-M). We also created a non-targeting, Rac2 *wild-type* control to test the effects of wild-type Rac2 (NT-Rac2^WT^-CAR-M). Lastly, an NG2-targeting wild-type Rac2 control was created to test for the effects of overexpressing Rac2 protein in addition to target binding without the hyperactivating variant (NG2-Rac2^WT^-CAR-M). We performed CAR-M-mediated engulfment assays as described above and found that NG2-Rac2^E62K^-CAR-MS exhibited significantly higher levels of GBM^U87^ engulfment than control macrophages (**Figure 4B; Figure S4A**). Interestingly, NG2-Rac2^E62K^-CAR-Ms exhibited similar levels of GBM^U87^ engulfment as NG2-Rac2^WT^-CAR-Ms (**Figure 4B; Figure S4A**), suggesting that overexpressing either Rac2^WT^ or Rac2^E62K^ enhances NG2-CAR-M-mediated phagocytosis over control macrophages. We then quantified GBM^U87^ spheroid growth in suspension over time, and found that spheroid growth was most inhibited with NG2-Rac2^E62K^-CAR-Ms or NG2-Rac2^WT^-CAR-Ms, compared to the original NG2-CAR-M and all other control CAR-Ms (**Figure 4C,D; Figure S4B**). Together, these data suggest that enhancing NG2-CAR-M phagocytosis capacity leads to enhanced inhibition of GBM spheroid growth.

**Figure 4:**
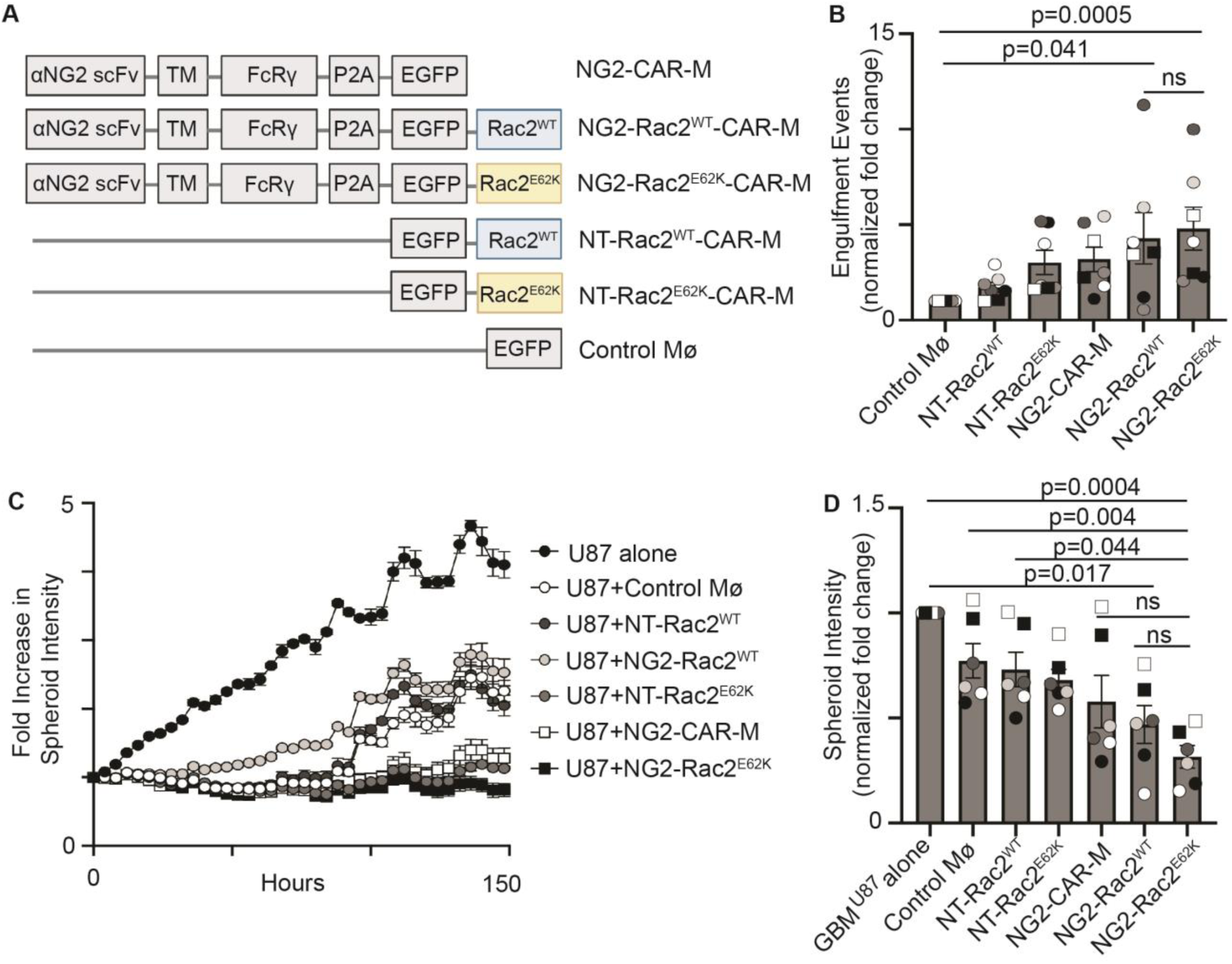
NG2-targeting CAR-Ms containing the Rac2^E62K^ hyperphagocytic signaling cassette exhibit higher rates engulfment and GBM spheroid growth control. A. Schematic of NG2-targeting CAR-Ms containing the Rac2^E62K^ signaling cassette or Rac2^WT^ and all associated non-targeting controls. B. Quantification of EGFP+ CAR-M-mediated engulfment of RFP+ GBM^U87^ cells by flow cytometry after 24 hours. n=7 independent PBMC donors (biological replicates), each with 3 technical replicates. The average of technical replicates for each PBMC donor is graphed and normalized to control macrophages. mean +/- SEM, Friedman’s test with Dunn’s multiple comparison. C. Representative images and graph of GBM^U87^ spheroid growth over time when cultured alone or with each CAR-M shown. Data are presented as fold changes in U87 RFP fluorescence intensity at each time point relative to the initial time point. D. Quantification of data in (C) at day 10 across conditions (n=6 independent PBMC donors (biological replicates), each with 6 technical replicates for every condition). Data are shown as fold increase in spheroid fluorescence intensity for each biological replicate and normalized to the GBM^U87^ alone control. mean +/- SD, Friedman’s test with Dunn’s multiple comparison.

Our data using the human immortalized GBM cell line, U87-MG, show that NG2-targeting CAR-Ms engulf human GBM cells and inhibit GBM spheroid growth. We next wanted to test whether CAR-Ms inhibited GBM engulfment and spheroid growth using patient-derived lines, such as the MD Anderson glioma stem cell (MDA-GSC) lines, that better capture the tumor heterogeneity and behavior observed in the clinical setting ^79^. We first identified glioma stem cell lines with high or low/no NG2 expression. We found that 92.9 +/- 7.6% of MDA-GSC 262 cells expressed surface NG2, as detected with the NG2^(9.2.27)^ antibody (**Figure 5A**) and 95.2 +/- 3.5% of MDA- GSC 262 cells expressed surface NG2 that is detected by the NG2^763.74^ antibody (**Figure 5B**). We identified another glioma stem cell line, MDA-GSC 8-11, that exhibited low to no expression of surface NG2 (**Figure 5A,B**). We generated MDA-GSC 262 and MDA-GSC 8-11 lines stably expressing luciferase/mCherry (herein referred to as GBM^GSC262^ and GBM^GSC811^) and asked whether NG2-CAR-Ms engulf glioma stem cells and inhibit their growth. For these studies, since we observed no significant differences in GBM^U87^ engulfment or spheroid inhibition between CAR-Ms expressing Rac2^WT^ versus Rac2^E62K^ (**Figure 4B-D**), we proceeded only with CARs containing Rac2^E62K^. We also made an additional modification to the NG2-CARs by “humanizing” the CAR. Instead of using the *murine* FcR gamma signaling domain as was used in our previous assays and in previous reports ^70,72,78^, we used the *human* FcR gamma signaling domain (**Figure S5A**). Concomitant with this change, we used the FcR gamma transmembrane sequence rather than the CD8 transmembrane sequence (**Figure S5A**). We found that NG2-CAR-Ms with the human FcR gamma signaling domain and transmembrane sequences showed higher rates of GBM engulfment compared to the NG2-CAR-M with murine sequences (**Figure S5B**). The NG2-CAR-Ms with the human FcR gamma signaling domain and transmembrane region will herein be referred to as *h*NG2-CAR-Ms. With these humanized CAR-Ms, we first determined whether hNG2-Rac2^E62K^-CAR-Ms exhibit differences in engulfment compared to control CAR-Ms.

**Figure 5:**
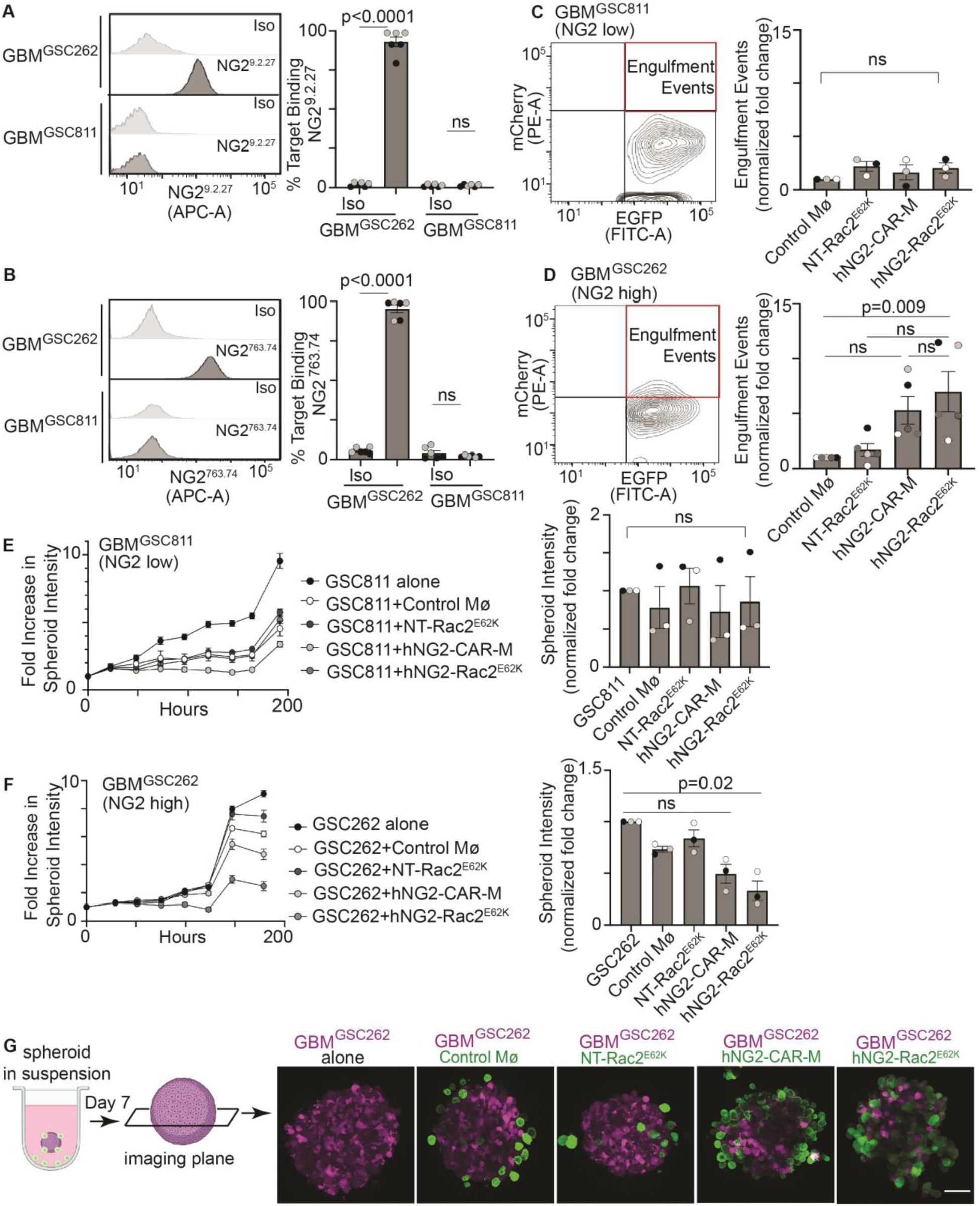
Humanized NG2-Rac2^E62K^ CAR-Ms engulf patient-derived glioblastoma stem cells and inhibit spheroid growth. A. Representative flow cytometry histogram (left) and quantification (right) of GBM^GSC262^ or GBM^GSC811^ cells surface stained with the NG2^9.2.27^ antibody versus an isotype control (Iso). B. Representative flow cytometry histogram (left) and quantification (right) of GBM^GSC262^ or GBM^GSC811^ cells surface stained with the NG2^763.74^ antibody versus an isotype control (Iso). For A and B, N=2 biological replicates. 2-way ANOVA with Tukey’s multiple comparison test. C. Representative flow cytometry graph (left) and quantification (right) of EGFP+ CAR-M-mediated engulfment of mCherry+ GBM^GSC811^ cells by flow cytometry after 24 hours. n=3 independent PBMC donors (biological replicates), each with 2 technical replicates. The average of technical replicates for each PBMC donor is graphed and normalized to control macrophages. mean +/- SEM, Friedman’s test with Dunn’s multiple comparison. D. Representative flow cytometry graph (left) and quantification (right) of EGFP+ CAR-M-mediated engulfment of mCherry+ GBM^GSC262^ cells by flow cytometry after 24 hours. n=5 independent PBMC donors (biological replicates), each with 2 technical replicates. The average of technical replicates for each PBMC donor is graphed and normalized to control macrophages. mean +/- SEM, Friedman’s test with Dunn’s multiple comparison. E. (left) Representative graph of GBM^GSC811^ spheroid growth over time when cultured alone or with each CAR-M shown. Data are presented as fold fluorescence intensity changes at each time point relative to the initial time point. (right) Quantification of data on left at day 10 across conditions (n=3 independent PBMC donors (biological replicates), each with 6 technical replicates for every condition). The average of technical replicates for each PBMC donor is graphed and normalized to GBM^GSC811^ alone control. mean +/- SEM, Friedman’s test with Dunn’s multiple comparison. F. (left) Representative graph of GBM^GSC262^ spheroid growth over time when cultured alone or with each CAR-M shown. Data are presented as fold fluorescence intensity changes at each time point relative to the initial time point. (right) Quantification of data on left at day 10 across conditions (n=3 independent PBMC donors (biological replicates), each with 6 technical replicates for every condition). The average of technical replicates for each PBMC donor is graphed and normalized to GBM^GSC262^ alone control. mean +/- SEM, Friedman’s test with Dunn’s multiple comparison. G. Representative images of GBM^GSC262^ spheroids that were cultured with CAR-Ms for 7 days in suspension and then removed from suspension for imaging. The schematic to the left shows the imaging plane through the center of the spheroid. Scale bar = 50 μm.

With GBM^GSC811^ (NG2^low^) cells, we found no significant difference in NG2-CAR-M-mediated engulfment across all CAR-Ms tested, including with hNG2-Rac2^E62K^-CAR-M, as expected (**Figure 5C; Figure S6A**). However, with GBM^GSC262^ (NG2^high^) cells, the only CAR-M that consistently and statistically significantly increased engulfment compared to control macrophages was hNG2-Rac2^E62K^-CAR-Ms (**Figure 5D; Figure S6B**). We then quantified glioma stem cell spheroid growth over 7 days. Again, as expected, GBM^GSC811^ (NG2 ^low^) cells did not exhibit significant differences in spheroid growth in the presence of any of the CAR-Ms (**Figure 5E; Figure S6C**). However, consistent with the GBM^GSC262^ (NG2^high^) engulfment results (**Figure 5F**), hNG2-Rac2^E62K^-CAR-M was the only CAR-M that significantly inhibited GBM^GSC262^ (NG2^high^) spheroid growth compared to GBM^GSC262^ (NG2^high^) spheroids alone (**Figure 5F; Figure S6D**). At day 7 of spheroid growth, the spheroids were removed from suspension and then visualized with higher resolution. We found that when we imaged the center of each spheroid, control macrophages and non-targeting CAR-Ms were not present at high numbers; whereas hNG2-Rac2^E62K^-CAR-Ms, and to a lesser extent, hNG2-CAR-Ms, exhibited high spheroid infiltration (**Figure 5G**), which again supported the spheroid growth and engulfment results.

Together, these results suggest that NG2-targeting CAR-Ms inhibit the growth of NG2-expressing patient-derived glioma stem cell spheroids.

We next sought to better understand how NG2-targeting CAR-Ms regulate the behavior of GBM^GSC262^ (NG2^high^) cells. We moved forward with only the hNG2-Rac2^E62K^-CAR-Ms for these studies since the hNG2-Rac2^E62K^-CAR-Ms showed significantly higher rates of GBM engulfment and spheroid control compared to control macrophages and GBM^GSC262^ spheroids alone, respectively (**Figure 5D,F**). We first evaluated GBM invasion of GBM^GSC262^ (NG2-high), GBM^GSC811^ (NG2-low), or GBM^U87^ (NG2-high cell line) cells. We embedded spheroids made of each of these lines in a 3D matrix and monitored their growth and invasion at 4 days post-embedding. We found that spheroids made with GBM^GSC262^ cells (NG2-high) or GBM^U87^ cells (NG2-high cell line) exhibited local invasion into the matrix at day 4, similar to the known invasive behavior of GBM in vivo, whereas spheroids made with GBM^GSC811^ cells (NG2-low) did not locally invade (**Figure S7**). We, therefore, used GBM^GSC262^ (NG2-high) cells to determine whether CAR-Ms regulated GBM invasion using time-lapse imaging. We generated GBM^GSC262^-Luc/mCherry spheroids and embedded them in a 3D matrix, with EGFP+ CAR-Ms distributed throughout the matrix (**Figure 6A**). We first asked whether CAR-Ms infiltrated GBM^GSC262^ spheroids. For better CAR-M visualization, we mosaically labeled spheroids with 10% Lck-mScarlet-expressing GBM^GSC262^ cells and 90% unlabeled GBM^GSC262^ cells. After 2 days of culture with hNG2-Rac2^E62K^-CAR-Ms, we imaged the center of the spheroid (**Figure 6B**). Using brightfield to outline the periphery of the GBM^GSC262^ spheroid (**Figure 6B**, white dotted outline), we found that EGFP-positive hNG2-Rac2^E62K^-CAR-Ms were present inside the spheroid, showing effective CAR-M infiltration when embedded in a 3D matrix (**Figure 6B**, arrows).

**Figure 6:**
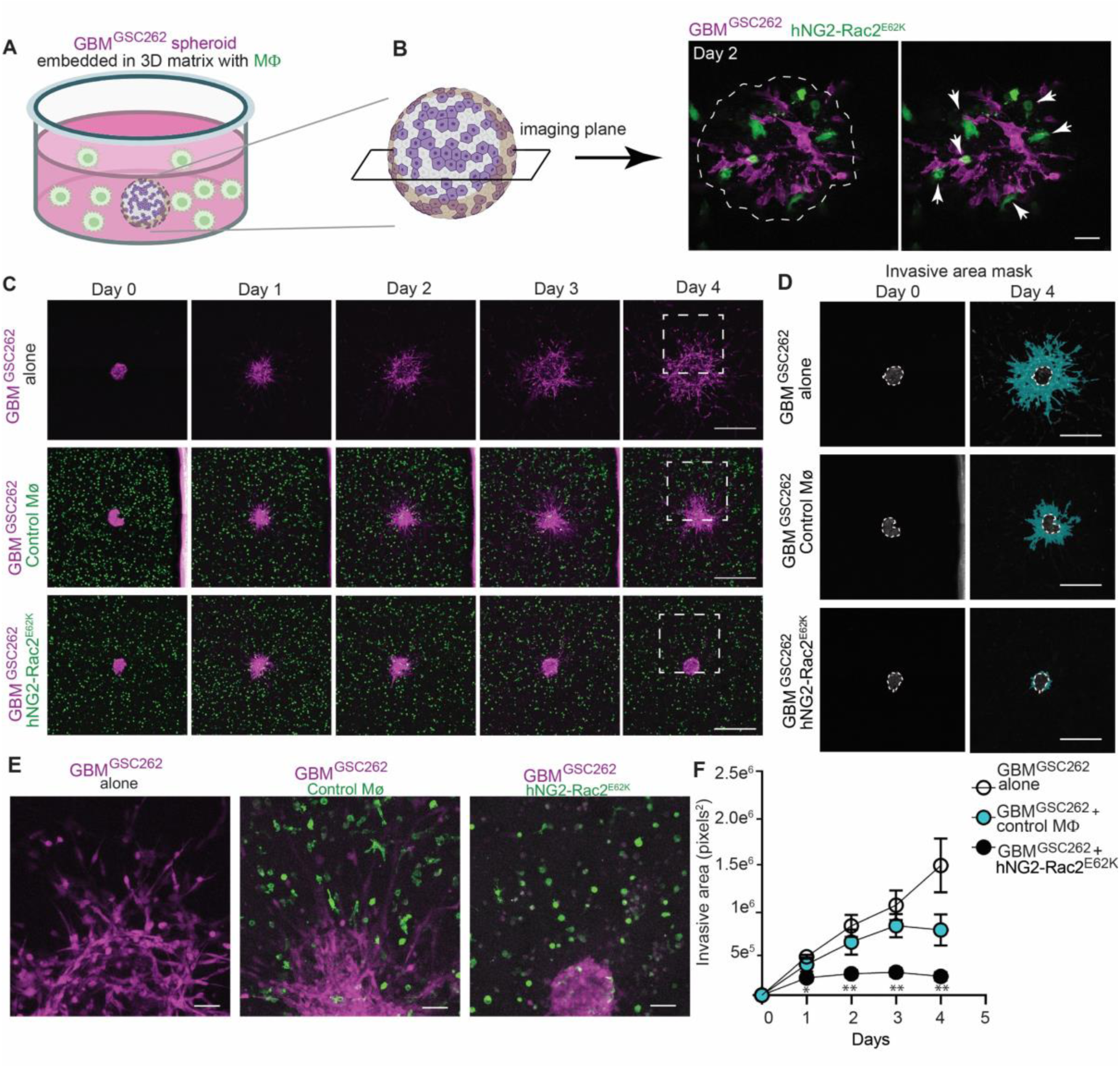
Humanized NG2-Rac2^E62K^ CAR-Ms inhibit 3D invasion of patient-derived glioblastoma stem cells. A. Schematic of mCherry+ GBM^GSC262^ spheroids embedded in a 3D matrix (50% Matrigel) with EGFP+ CAR-Ms. B. Schematic of 3D embedded GBM^GSC262^ spheroid, and the imaging plane used to acquire images to visualize CAR-M infiltration into the spheroid. GBM^GSC262^ spheroids were comprised of 10% mCherry-positive GBM^GSC262^ cells and 90% unlabelled GBM^GSC262^ cells to better reveal EGFP+ CAR-M infiltration. The spheroid is outlined in white, and the arrowheads point to hNG2-Rac2^E62K^ CAR-Ms that have infiltrated the spheroid. Scale bar = 50 μm. C. Representative images of GBM^GSC262^ spheroids (magenta) embedded in 3D matrices over 4 days when embedded alone (top panels), or in the presence of control macrophages (middle panels, green) or hNG2-Rac2^E62K^ CAR-Ms (bottom panels, green). Scale bar = 500 μm. D. Representative images of GBM^GSC262^ spheroid channel only at day 0 and day 4, with the spheroid at time zero outlined in white and the invasive area pseudocolored in teal. Scale bar = 50 μm. E. Zoom of outlined squares in C. F. Quantification of invasive area over time (n=5 spheroids across n=3 PBMC donors (biological replicates)). mean +/- SEM, Kruskal-Wallis test at each time point comparing the invasive area of GBM^GSC262^ spheroids treated with hNG2-Rac2^E62K^ CAR-Ms compared to GBM^GSC262^ spheroids alone. *p<0.05 **p<0.01

We next tested whether CAR-Ms regulate GBM^GSC262^ cell invasion from the spheroid and into the matrix. We made spheroids with 100% of mCherry-expressing GBM^GSC262^ cells and observed that, as we described above (**Figure S7**), GBM^GSC262^ cells exhibited high levels of local invasion into the matrix over 4 days (**Figure 6C**, top panels). In the presence of hNG2-Rac2^E62K^-CAR-Ms, GBM^GSC262^ cell invasion was dramatically reduced compared to the control macrophage condition and GBM^GSC262^ cells alone (**Figure 6C-F**). To quantify local invasion, we measured spheroid spread over time by quantifying the area of invaded cells and subtracting the spheroid size at time zero (**Figure 6D**, invasive area pseudocolored in teal; **Figure 6F**).

These data show that local spheroid invasion was decreased by ∼85% at day 4 in the presence of hNG2-Rac2^E62K^-CAR-Ms compared to GBM^GSC262^ spheroids alone.

To determine whether CAR-Ms *prevent* GBM invasion into the matrix or *engulf* GBM cells that locally invade into the matrix, or both, we examined CAR-M/GBM interactions in greater detail using high-resolution time-lapse imaging (**Figure 7**). We focused on days 2-3 after spheroid embedding because this is the time point at which differences in GBM^GSC262^ local invasion in the matrix between control and hNG2-Rac2^E62K^-CAR-M-treated GBM spheroids become apparent. We also made spheroids using a combination of H2B-mCherry and Lck-mScarlet-expressing GBM^GSC262^ cells to better track GBM^GSC262^ cells with the H2B nuclear label and identify spheroid margins with the Lck-mScarlet membrane localization. We found that in the presence of control macrophages, H2B-mcherry-expressing GBM^GSC262^ cells move away from the spheroid and invade into the 3D matrix (**Figure 7A; Movies S1-S3**). However, surprisingly, in the presence of hNG2-Rac2^E62K^-CAR-Ms, H2B-mcherry-expressing GBM^GSC262^ cells did not appear to invade into the matrix; rather, GBM^GSC262^ cells moved within the spheroid but did not translocate away from the spheroid (**Figure 7B**). These results suggest that hNG2-Rac2^E62K^-CAR-Ms prevent GBM^GSC262^ cell invasion from the spheroid.

**Figure 7:**
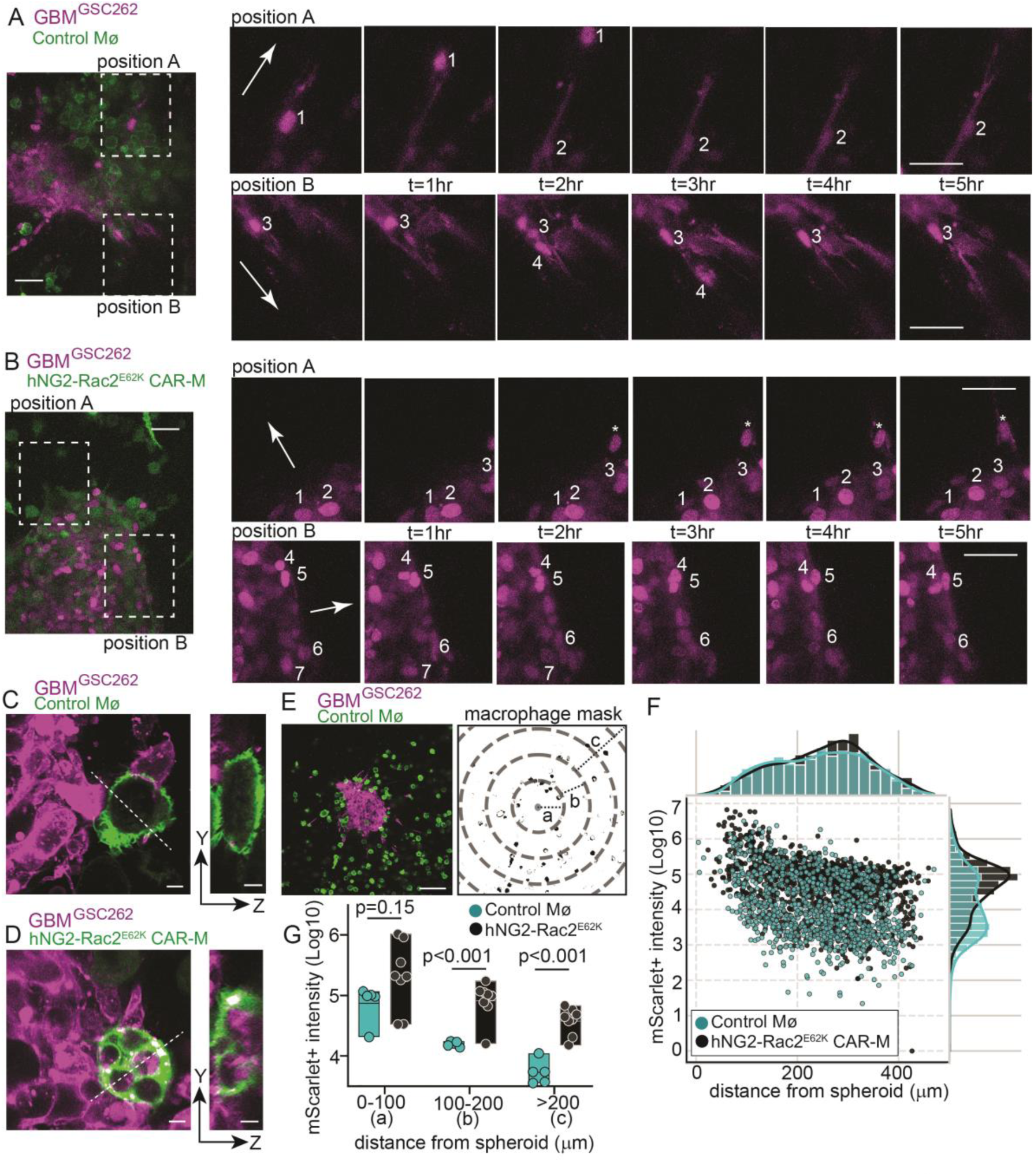
Patient-derived glioblastoma stem cells are engulfed by NG2-Rac2^E62K^ CAR-Ms and do not locally invade in the presence of NG2-Rac2^E62K^ CAR-Ms. A. H2B-mCherry/mCherry-expressing GBM^GSC262^ spheroids (magenta) embedded in a 3D matrix with EGFP-positive control macrophages (green). Outlined boxes indicate zoomed positions on the right, showing mCherry-positive GBM^GSC262^ cells only over time. Arrows in the first panel indicate the direction away from the spheroid, and individual H2B-mCherry GBM^GSC262^ cell nuclei are numbered. Time is indicated in hours. B. H2B-mCherry/Lck-mScarlet-expressing GBM^GSC262^ spheroids (magenta) embedded in a 3D matrix with EGFP-positive hNG2-Rac2^E62K^ CAR-Ms (green). Outlined boxes indicate zoomed positions on the right, showing mCherry-positive GBM^GSC262^ cells only over time. Arrows in the first panel indicate the direction away from the spheroid, and individual H2B-mCherry GBM^GSC262^ cell nuclei are numbered. Asterisk marks an H2B-mCherry GBM^GSC262^ cell that is inside of an hNG2-Rac2^E62K^ CAR-M. Time is indicated in hours. For A and B, scale bars = 50 μm. C. Representative image of an EGFP+ control macrophage (green) at the edge of a GBM^GSC262^ spheroid (magenta) in a 3D matrix, exhibiting no engulfment. Y-Z analysis of the white dotted line shows no internalized Lck-mScarlet+ GBM^GSC262^ fragments. D. Representative image of an hNG2-Rac2^E62K^ CAR-M (green) at the edge of a GBM^GSC262^ spheroid (magenta) in a 3D matrix, exhibiting GBM^GSC262^ engulfment. Y-Z analysis of the white dotted line shows several internalized Lck-mScarlet+ GBM^GSC262^ fragments. For C and D, scale bars = 10 μm. E. (Left) Representative image of Lck-mScarlet+ GBM^GSC262^ spheroid (magenta) with control EGFP+ macrophages (green) in a 3D matrix. Scale bar = 100 μm. (Right) Example of thresholding to create a mask for EGFP+ macrophages to then quantify Lck-mScarlet+ fluorescence intensity within the masked macrophages. Concentric rings each reflect intervals of 100 μm, annotated as “a” (0-100 μm), “b” (100 μm-200 μm), and “c” (>200 μm) from the center of the spheroid. F. Quantification of Lck-mScarlet+ fluorescence intensity in control macrophages (blue) versus hNG2-Rac2^E62K^ CAR-Ms (black) across the Y-axis (histogram to the right) and quantification of the position of control macrophages (blue) versus hNG2-Rac2^E62K^ CAR-Ms (black) relative to the spheroid across the X-axis (histogram at the top). Each dot represents an individual macrophage from control macrophages (n=1069) and hNG2-Rac2^E62K^ CAR-Ms (n=1166). G. Quantification of the median Lck-mScarlet+ fluorescence intensity in control macrophages (blue) versus hNG2-Rac2^E62K^ CAR-Ms (black) for each spheroid: Control macrophages (n=5 spheroids); hNG2-Rac2^E62K^ CAR-Ms (n=8 spheroids).

We then compared the GBM^GSC262^ cell engulfment capacity of hNG2-Rac2^E62K^-CAR-Ms versus control macrophages in the 3D matrix at the same time point as above (2-3 days). We first observed that hNG2-Rac2^E62K^-CAR-Ms adjacent to the spheroid exhibited large amounts of internalized Lck-mScarlet+ GBM^GSC262^ cell fragments compared to control macrophages (**Figure 7C, D**). To quantify this engulfment, we created masks on the EGFP+ macrophages. We then quantified the fluorescence intensity of Lck-mScarlet+ GBM^GSC262^ cell fragments within those macrophages as a function of distance from the spheroid, with each concentric ring as 100μm from the center of the spheroid (**Figure 7E**). We first found that the distribution of hNG2-Rac2^E62K^-CAR-Ms versus control macrophages from the spheroid was similar (**Figure 7F**, X-axis histogram). However, we found that hNG2-Rac2^E62K^-CAR-Ms contained significantly higher levels of Lck-mScarlet+ GBM^GSC262^ cell fragments compared to control macrophages (**Figure 7F**, Y-axis histogram). We then quantified the fluorescence intensity of Lck-mScarlet+ GBM^GSC262^ cell fragments in macrophages that are within the spheroid (0-100μm, “a” in Figure 7E), adjacent to the spheroid (100-200μm, “b” in Figure 7E), or at a further distance from the spheroid (>200μm, “c” in Figure 7E). We found that as the distance from the spheroid increased, the fluorescent intensity of Lck-mScarlet+ GBM^GSC262^ cell fragments in both hNG2-Rac2^E62K^-CAR-Ms and control macrophages decreased (**Figure 7G**). There was no statistically significant difference in the level of Lck-mScarlet+ GBM^GSC262^ cell fragments between hNG2-Rac2^E62K^-CAR-Ms and control macrophages inside the spheroid (0-100μm); however, at areas adjacent to the spheroid (100-200μm) and at distances further away (>200μm), hNG2-Rac2^E62K^-CAR-Ms contained significantly higher levels Lck-mScarlet+ GBM^GSC262^ cell fragments compared to control macrophages (**Figure 7G**). Together, these data suggest that hNG2-Rac2^E62K^-CAR-Ms both engulf GBM^GSC262^ cells and inhibit their local invasion.

## DISCUSSION

A small number of studies show that CAR-Ms designed to recognize glioblastoma antigens cause tumor regression and reprogramming of the immune system ^54-57^. These studies have been enormously valuable and highlight the promise for CAR-Ms as a therapeutic, but what still remains unknown is how CAR-Ms dynamically interact with GBM cells to mediate this response. In this work, we fill this gap in knowledge by directly visualizing CAR-M and GBM interactions at high resolution and over time. As expected, we found that NG2-targeting CAR-Ms engulf GBM cells. However, unexpectedly, we found that NG2-targeting CAR-Ms inhibit GBM local invasion, and that this inhibition occurs through both GBM engulfment and inhibition of GBM invasion from the spheroid. This work provides the foundation for future experiments delving into how CAR-Ms may prevent GBM local invasion from the tumor, and how we can leverage these findings to develop targeted therapeutics.

A key question that arises from our findings is how CAR-Ms inhibit GBM local invasion. This result is particularly interesting in light of numerous papers showing that macrophages *promote* GBM invasion. In these examples, these macrophages are tumor-associated macrophages and significantly enhance the invasive spread of GBM cells ^19–22,24^. In the case of CAR-Ms, we show that NG2-targeting CAR-Ms exhibit superior GBM engulfment than control macrophages in 3D, suggesting that targeted GBM engulfment contributes to inhibition of GBM invasion (**Figure 7C-G**). However, strikingly, in the presence of NG2-targeting CAR-Ms, but not control macrophages, GBM cells did not appear to invade into the matrix (**Figure 7A, B**), suggesting a potential engulfment-independent function for CAR-Ms. Previous studies have examined changes to macrophage differentiation and secretion profiles after engulfing a target and shown that macrophages alter their differentiation state and secrete signals to promote wound healing and tissue remodeling ^54,56,80–83^. Thus, we initially hypothesized that NG2-targeting CAR-Ms might also shift their macrophage differentiation toward a more anti-tumor-like state, and secrete specific signals to prevent GBM invasion. To test this hypothesis, we analyzed the macrophage differentiation status of CAR-Ms. We co-cultured GBM^GSC262^ (NG2^high^) cells with hNG2-Rac2^E62K^-CAR-Ms or control CAR-Ms and quantified the expression of pro-inflammatory (ie. “anti-tumor”) markers on the macrophages after 3 days of coculture. We selected 3 days of co-culture, as this is the time point (2-3 days) at which we first observe differences in GBM invasion in the presence of NG2-CAR-Ms compared to control macrophages. We found no difference in the expression profiles of these pro-inflammatory/anti-tumor markers in NG2-CAR-Ms compared to any of the controls (**Figure S8A-C**), indicating that our data are inconsistent with this hypothesis and suggesting that NG2-CAR-Ms do not exhibit significant shifts in macrophage differentiation state compared to control macrophages. These results suggest that although NG2-CAR-Ms do not change their expression of anti-tumor markers, they may still be secreting factors that limit GBM invasion. It is also possible that GBM cells that have been eaten, but not killed, by NG2-targeting CAR-Ms secrete signals to neighboring GBM cells. In support of this hypothesis, target cell engulfment within a tissue is regulated by the behavior of surrounding cells, which change shape and send signals to neighboring cells to maintain homeostasis ^84–86^. Thus, GBM cells that have been eaten may send signals to, or regulate the mechanical tension of, nearby GBM cells to downregulate global invasion programs. GBM cells that have been eaten by NG2-targeting CAR-Ms may also downregulate their own invasion programs, as previous work has shown that tuning cell death signaling potently inhibits cell migration ^87^.

An alternative explanation for the inhibition of GBM local invasion is through changes in the extracellular matrix. Macrophages remodel the extracellular matrix directly by secreting a broad array of matrix metalloproteinases ^88,89^ and/or by regulating matrix stiffness through matrix cross-linking and deposition ^90–92^. Previous experiments have shown that tumor-associated macrophage-dependent remodeling of the extracellular matrix increases cancer cell invasion ^92^. Macrophages also secrete signals that regulate the expression of matrix metalloproteases in cancer cells ^93–95^. Thus, NG2-targeting macrophages may remodel the matrix either directly or indirectly, thereby reducing GBM invasion. Future experiments dissecting whether signals from neighboring GBM cells or CAR-Ms are altered in the presence of NG2-targeting CAR-Ms and probing extracellular matrix organization and stiffness will help shed light on each of these processes.

The poor prognosis of patients with GBM is due to the ability of GBM cells to leave the primary tumor and invade throughout the brain, resulting in microscopic residual disease even after gross total tumor resection. This residual disease leads to high rates of recurrence. Thus, inhibiting GBM invasion, survival, and growth are key steps to inhibit in the clinic. For CAR-Ms to be an effective therapeutic strategy, a better understanding of whether CAR-Ms disseminate throughout the brain, and how CAR-Ms dynamically interact with GBM cells, is critical. Will targeted engulfment of microscopic disease by CAR-Ms be the key step, or CAR-M-mediated reprogramming of immune cells, or both? Future experiments using in vivo models to directly visualize CAR-M interactions with GBM tumors and with components of the immune system are valuable next steps.

## ACKNOWLEDGEMENTS AND FUNDING SOURCES

We would like to thank members of the Roh-Johnson lab for their constructive critiques on this work. We would also like to thank Dr. Frederick F. Lang (University of Texas MD Anderson Cancer Center) for sharing patient-derived glioma stem cell lines; Dr. Kaitlin Basham (University of Utah, Huntsman Cancer Institute) for sharing plasmids; and Dr. Keren Hilgendorf (University of Utah) for use of their equipment. This work was supported by the National Cancer Institute R37CA247994, the Huntsman Cancer Institute Translational Science Initiative Award 24-TSI-02, the University of Utah Ascender Grant U-8006, and a gift from CaLycia Bioscience (to MRJ);

Neurosurgery Research Education Foundation Grant 10067440 (to EK); NIH/NCI T32CA265782 (to SRSV); S.R. Lowry Endowed Chair in the Department of Neurosurgery, Ty Louis Cambel Foundation, and the Tate Family (to SHC); NIH Director’s Pioneer Award 5DP1CA300850 (to DJM); Huntsman Cancer Institute Cancer Center Support Grant P30CA040214 and the V Foundation V2024-002 (to MQR); Office Of The Director of the National Institutes of Health S10OD026959 and National Cancer Institute 5P30CA042014-24 (to Flow Cytometry Core).

## AUTHOR CONTRIBUTIONS

EK, SRSV, and DG conceptualized the project, designed and conducted experiments, analyzed and interpreted data, managed collaborations, and obtained funding. MRJ supervised and conceptualized the project, interpreted data, managed collaborations, wrote the manuscript, and obtained funding. JEM and GH conducted experiments and analyzed data. MQR, DJM, and SHC helped with data interpretation and obtained funding. GD, HC, AP, AKM, and MR provided resources. All authors provided edits for the manuscript.

## ETHICS APPROVAL AND CONSENT TO PARTICIPATE

The blood donors for monocyte isolation were obtained from a commercial pathology testing vendor (ARUP Laboratories) and were de-identified. Therefore, we do not have information regarding age, sex, or consent, nor do we require IRB approval.

## DATA AVAILABILITY STATEMENT

The data generated in this study are available within the article and its corresponding supplementary data files.

## REFERENCES

1. Grochans, S., Cybulska, A.M., Siminska, D., Korbecki, J., Kojder, K., Chlubek, D., and Baranowska-Bosiacka, I. (2022). Epidemiology of Glioblastoma Multiforme-Literature Review. Cancers (Basel) 14. 10.3390/cancers14102412.

2. Stupp, R., Taillibert, S., Kanner, A., Read, W., Steinberg, D., Lhermitte, B., Toms, S., Idbaih, A., Ahluwalia, M.S., Fink, K., et al. (2017). Effect of Tumor-Treating Fields Plus Maintenance Temozolomide vs Maintenance Temozolomide Alone on Survival in Patients With Glioblastoma: A Randomized Clinical Trial. JAMA 318, 2306–2316. 10.1001/jama.2017.18718.

3. Boumahdi, S., and de Sauvage, F.J. (2020). The great escape: tumour cell plasticity in resistance to targeted therapy. Nat Rev Drug Discov 19, 39–56. 10.1038/s41573-019-0044-1.

4. Neftel, C., Laffy, J., Filbin, M.G., Hara, T., Shore, M.E., Rahme, G.J., Richman, A.R., Silverbush, D., Shaw, M.L., Hebert, C.M., et al. (2019). An Integrative Model of Cellular States, Plasticity, and Genetics for Glioblastoma. Cell 178, 835–849 e821. 10.1016/j.cell.2019.06.024.

5. Sharma, P., Aaroe, A., Liang, J., and Puduvalli, V.K. (2023). Tumor microenvironment in glioblastoma: Current and emerging concepts. Neurooncol Adv 5, vdad009. 10.1093/noajnl/vdad009.

6. Martinez-Lage, M., Lynch, T.M., Bi, Y., Cocito, C., Way, G.P., Pal, S., Haller, J., Yan, R.E., Ziober, A., Nguyen, A., et al. (2019). Immune landscapes associated with different glioblastoma molecular subtypes. Acta Neuropathol Commun 7, 203. 10.1186/s40478-019-0803-6.

7. Anderson, N.M., and Simon, M.C. (2020). The tumor microenvironment. Curr Biol 30, R921–R925. 10.1016/j.cub.2020.06.081.

8. Schreiber, R.D., Old, L.J., and Smyth, M.J. (2011). Cancer immunoediting: integrating immunity’s roles in cancer suppression and promotion. Science 331, 1565–1570. 10.1126/science.1203486.

9. Qian, J., Luo, F., Yang, J., Liu, J., Liu, R., Wang, L., Wang, C., Deng, Y., Lu, Z., Wang, Y., et al. (2018). TLR2 Promotes Glioma Immune Evasion by Downregulating MHC Class II Molecules in Microglia. Cancer Immunol Res 6, 1220–1233. 10.1158/2326-6066.CIR-18-0020.

10. Zhao, J., Chen, A.X., Gartrell, R.D., Silverman, A.M., Aparicio, L., Chu, T., Bordbar, D., Shan, D., Samanamud, J., Mahajan, A., et al. (2019). Immune and genomic correlates of response to anti-PD-1 immunotherapy in glioblastoma. Nat Med 25, 462–469. 10.1038/s41591-019-0349-y.

11. O’Rourke, D.M., Nasrallah, M.P., Desai, A., Melenhorst, J.J., Mansfield, K., Morrissette, J.J.D., Martinez-Lage, M., Brem, S., Maloney, E., Shen, A., et al. (2017). A single dose of peripherally infused EGFRvIII-directed CAR T cells mediates antigen loss and induces adaptive resistance in patients with recurrent glioblastoma. Sci Transl Med 9. 10.1126/scitranslmed.aaa0984.

12. Xie, Y.J., Dougan, M., Jailkhani, N., Ingram, J., Fang, T., Kummer, L., Momin, N., Pishesha, N., Rickelt, S., Hynes, R.O., and Ploegh, H. (2019). Nanobody-based CAR T cells that target the tumor microenvironment inhibit the growth of solid tumors in immunocompetent mice. Proc Natl Acad Sci U S A 116, 7624–7631. 10.1073/pnas.1817147116.

13. Ma, J., Chen, C.C., and Li, M. (2021). Macrophages/Microglia in the Glioblastoma Tumor Microenvironment. Int J Mol Sci 22. 10.3390/ijms22115775.

14. Andersen, R.S., Anand, A., Harwood, D.S.L., and Kristensen, B.W. (2021). Tumor-Associated Microglia and Macrophages in the Glioblastoma Microenvironment and Their Implications for Therapy. Cancers (Basel) 13. 10.3390/cancers13174255.

15. Hambardzumyan, D., Gutmann, D.H., and Kettenmann, H. (2016). The role of microglia and macrophages in glioma maintenance and progression. Nat Neurosci 19, 20–27. 10.1038/nn.4185.

16. Mitsdoerffer, M., Aly, L., Barz, M., Engleitner, T., Sie, C., Delbridge, C., Lepennetier, G., Ollinger, R., Pfaller, M., Wiestler, B., et al. (2022). The glioblastoma multiforme tumor site promotes the commitment of tumor-infiltrating lymphocytes to the T(H)17 lineage in humans. Proc Natl Acad Sci U S A 119, e2206208119. 10.1073/pnas.2206208119.

17. Guo, X., Pan, Y., and Gutmann, D.H. (2019). Genetic and genomic alterations differentially dictate low-grade glioma growth through cancer stem cell-specific chemokine recruitment of T cells and microglia. Neuro Oncol 21, 1250–1262. 10.1093/neuonc/noz080.

18. Held-Feindt, J., Hattermann, K., Muerkoster, S.S., Wedderkopp, H., Knerlich-Lukoschus, F., Ungefroren, H., Mehdorn, H.M., and Mentlein, R. (2010). CX3CR1 promotes recruitment of human glioma-infiltrating microglia/macrophages (GIMs). Exp Cell Res 316, 1553–1566. 10.1016/j.yexcr.2010.02.018.

19. Bettinger, I., Thanos, S., and Paulus, W. (2002). Microglia promote glioma migration. Acta Neuropathol 103, 351–355. 10.1007/s00401-001-0472-x.

20. Liu, Z., Kuang, W., Zhou, Q., and Zhang, Y. (2018). TGF-beta1 secreted by M2 phenotype macrophages enhances the stemness and migration of glioma cells via the SMAD2/3 signalling pathway. Int J Mol Med 42, 3395–3403. 10.3892/ijmm.2018.3923.

21. Markovic, D.S., Glass, R., Synowitz, M., Rooijen, N., and Kettenmann, H. (2005). Microglia stimulate the invasiveness of glioma cells by increasing the activity of metalloprotease-2. J Neuropathol Exp Neurol 64, 754–762. 10.1097/01.jnen.0000178445.33972.a9.

22. Markovic, D.S., Vinnakota, K., Chirasani, S., Synowitz, M., Raguet, H., Stock, K., Sliwa, M., Lehmann, S., Kalin, R., van Rooijen, N., et al. (2009). Gliomas induce and exploit microglial MT1-MMP expression for tumor expansion. Proc Natl Acad Sci U S A 106, 12530–12535. 10.1073/pnas.0804273106.

23. Wu, A., Wei, J., Kong, L.Y., Wang, Y., Priebe, W., Qiao, W., Sawaya, R., and Heimberger, A.B. (2010). Glioma cancer stem cells induce immunosuppressive macrophages/microglia. Neuro Oncol 12, 1113–1125. 10.1093/neuonc/noq082.

24. Ye, X.Z., Xu, S.L., Xin, Y.H., Yu, S.C., Ping, Y.F., Chen, L., Xiao, H.L., Wang, B., Yi, L., Wang, Q.L., et al. (2012). Tumor-associated microglia/macrophages enhance the invasion of glioma stem-like cells via TGF-beta1 signaling pathway. J Immunol 189, 444–453. 10.4049/jimmunol.1103248.

25. He, X., Guo, Y., Yu, C., Zhang, H., and Wang, S. (2023). Epithelial-mesenchymal transition is the main way in which glioma-associated microglia/macrophages promote glioma progression. Front Immunol 14, 1097880. 10.3389/fimmu.2023.1097880.

26. Gjorgjevski, M., Hannen, R., Carl, B., Li, Y., Landmann, E., Buchholz, M., Bartsch, J.W., and Nimsky, C. (2019). Molecular profiling of the tumor microenvironment in glioblastoma patients: correlation of microglia/macrophage polarization state with metalloprotease expression profiles and survival. Biosci Rep 39. 10.1042/BSR20182361.

27. Sorensen, M.D., Dahlrot, R.H., Boldt, H.B., Hansen, S., and Kristensen, B.W. (2018). Tumour-associated microglia/macrophages predict poor prognosis in high-grade gliomas and correlate with an aggressive tumour subtype. Neuropathol Appl Neurobiol 44, 185–206. 10.1111/nan.12428.

28. Quail, D.F., and Joyce, J.A. (2017). The Microenvironmental Landscape of Brain Tumors. Cancer Cell 31, 326–341. 10.1016/j.ccell.2017.02.009.

29. Greenwald, A.C., Darnell, N.G., Hoefflin, R., Simkin, D., Mount, C.W., Gonzalez Castro, L.N., Harnik, Y., Dumont, S., Hirsch, D., Nomura, M., et al. (2024). Integrative spatial analysis reveals a multi-layered organization of glioblastoma. Cell 187, 2485–2501 e2426. 10.1016/j.cell.2024.03.029.

30. Gholamin, S., Mitra, S.S., Feroze, A.H., Liu, J., Kahn, S.A., Zhang, M., Esparza, R., Richard, C., Ramaswamy, V., Remke, M., et al. (2017). Disrupting the CD47-SIRPalpha anti-phagocytic axis by a humanized anti-CD47 antibody is an efficacious treatment for malignant pediatric brain tumors. Sci Transl Med 9. 10.1126/scitranslmed.aaf2968.

31. Krausgruber, T., Blazek, K., Smallie, T., Alzabin, S., Lockstone, H., Sahgal, N., Hussell, T., Feldmann, M., and Udalova, I.A. (2011). IRF5 promotes inflammatory macrophage polarization and TH1-TH17 responses. Nat Immunol 12, 231–238. 10.1038/ni.1990.

32. Advani, R., Flinn, I., Popplewell, L., Forero, A., Bartlett, N.L., Ghosh, N., Kline, J., Roschewski, M., LaCasce, A., Collins, G.P., et al. (2018). CD47 Blockade by Hu5F9-G4 and Rituximab in Non-Hodgkin’s Lymphoma. N Engl J Med 379, 1711–1721. 10.1056/NEJMoa1807315.

33. Kaneda, M.M., Messer, K.S., Ralainirina, N., Li, H., Leem, C.J., Gorjestani, S., Woo, G., Nguyen, A.V., Figueiredo, C.C., Foubert, P., et al. (2016). PI3Kgamma is a molecular switch that controls immune suppression. Nature 539, 437–442. 10.1038/nature19834.

34. Guerriero, J.L., Sotayo, A., Ponichtera, H.E., Castrillon, J.A., Pourzia, A.L., Schad, S., Johnson, S.F., Carrasco, R.D., Lazo, S., Bronson, R.T., et al. (2017). Class IIa HDAC inhibition reduces breast tumours and metastases through anti-tumour macrophages. Nature 543, 428–432. 10.1038/nature21409.

35. O’Connell, B.C., Hubbard, C., Zizlsperger, N., Fitzgerald, D., Kutok, J.L., Varner, J., Ilaria, R., Jr., Cobleigh, M.A., Juric, D., Tkaczuk, K.H.R., et al. (2024). Eganelisib combined with immune checkpoint inhibitor therapy and chemotherapy in frontline metastatic triple-negative breast cancer triggers macrophage reprogramming, immune activation and extracellular matrix reorganization in the tumor microenvironment. J Immunother Cancer 12. 10.1136/jitc-2024-009160.

36. Zhao, W., Zhang, Z., Xie, M., Ding, F., Zheng, X., Sun, S., and Du, J. (2025). Exploring tumor-associated macrophages in glioblastoma: from diversity to therapy. NPJ Precis Oncol 9, 126. 10.1038/s41698-025-00920-x.

37. Li, T.F., Li, K., Wang, C., Liu, X., Wen, Y., Xu, Y.H., Zhang, Q., Zhao, Q.Y., Shao, M., Li, Y.Z., et al. (2017). Harnessing the cross-talk between tumor cells and tumor-associated macrophages with a nano-drug for modulation of glioblastoma immune microenvironment. J Control Release 268, 128–146. 10.1016/j.jconrel.2017.10.024.

38. Stupp, R., Mason, W.P., van den Bent, M.J., Weller, M., Fisher, B., Taphoorn, M.J., Belanger, K., Brandes, A.A., Marosi, C., Bogdahn, U., et al. (2005). Radiotherapy plus concomitant and adjuvant temozolomide for glioblastoma. N Engl J Med 352, 987–996. 10.1056/NEJMoa043330.

39. Sadelain, M., Brentjens, R., and Riviere, I. (2013). The basic principles of chimeric antigen receptor design. Cancer Discov 3, 388–398. 10.1158/2159-8290.CD-12-0548.

40. Rafiq, S., Hackett, C.S., and Brentjens, R.J. (2020). Engineering strategies to overcome the current roadblocks in CAR T cell therapy. Nat Rev Clin Oncol 17, 147–167. 10.1038/s41571-019-0297-y.

41. Lim, W.A., and June, C.H. (2017). The Principles of Engineering Immune Cells to Treat Cancer. Cell 168, 724–740. 10.1016/j.cell.2017.01.016.

42. Fesnak, A.D., June, C.H., and Levine, B.L. (2016). Engineered T cells: the promise and challenges of cancer immunotherapy. Nat Rev Cancer 16, 566–581. 10.1038/nrc.2016.97.

43. Bagley, S.J., Binder, Z.A., Lamrani, L., Marinari, E., Desai, A.S., Nasrallah, M.P., Maloney, E., Brem, S., Lustig, R.A., Kurtz, G., et al. (2024). Repeated peripheral infusions of anti-EGFRvIII CAR T cells in combination with pembrolizumab show no efficacy in glioblastoma: a phase 1 trial. Nat Cancer 5, 517–531. 10.1038/s43018-023-00709-6.

44. Begley, S.L., O’Rourke, D.M., and Binder, Z.A. (2025). CAR T cell therapy for glioblastoma: A review of the first decade of clinical trials. Mol Ther 33, 2454–2461. 10.1016/j.ymthe.2025.03.004.

45. Brown, C.E., Hibbard, J.C., Alizadeh, D., Blanchard, M.S., Natri, H.M., Wang, D., Ostberg, J.R., Aguilar, B., Wagner, J.R., Paul, J.A., et al. (2024). Locoregional delivery of IL-13Ralpha2-targeting CAR-T cells in recurrent high-grade glioma: a phase 1 trial. Nat Med 30, 1001–1012. 10.1038/s41591-024-02875-1.

46. Choi, B.D., Gerstner, E.R., Frigault, M.J., Leick, M.B., Mount, C.W., Balaj, L., Nikiforow, S., Carter, B.S., Curry, W.T., Gallagher, K., and Maus, M.V. (2024). Intraventricular CARv3-TEAM-E T Cells in Recurrent Glioblastoma. N Engl J Med 390, 1290–1298. 10.1056/NEJMoa2314390.

47. Gallus, M., Young, J.S., Cook Quackenbush, S., Khasraw, M., de Groot, J., and Okada, H. (2025). Chimeric antigen receptor T-cell therapy in patients with malignant glioma-From neuroimmunology to clinical trial design considerations. Neuro Oncol 27, 352–368. 10.1093/neuonc/noae203.

48. Pant, A., and Lim, M. (2023). CAR-T Therapy in GBM: Current Challenges and Avenues for Improvement. Cancers (Basel) 15. 10.3390/cancers15041249.

49. Brown, C.E., Alizadeh, D., Starr, R., Weng, L., Wagner, J.R., Naranjo, A., Ostberg, J.R., Blanchard, M.S., Kilpatrick, J., Simpson, J., et al. (2016). Regression of Glioblastoma after Chimeric Antigen Receptor T-Cell Therapy. N Engl J Med 375, 2561–2569. 10.1056/NEJMoa1610497.

50. Park, S., Maus, M.V., and Choi, B.D. (2024). CAR-T cell therapy for the treatment of adult high-grade gliomas. NPJ Precis Oncol 8, 279. 10.1038/s41698-024-00753-0.

51. Huang, J., Liu, F., Li, C., Liang, X., Li, C., Liu, Y., Yi, Z., Zhang, L., Fu, S., and Zeng, Y. (2022). Role of CD47 in tumor immunity: a potential target for combination therapy. Sci Rep 12, 9803. 10.1038/s41598-022-13764-3.

52. Kim, D., Wang, J., Willingham, S.B., Martin, R., Wernig, G., and Weissman, I.L. (2012). Anti-CD47 antibodies promote phagocytosis and inhibit the growth of human myeloma cells. Leukemia 26, 2538–2545. 10.1038/leu.2012.141.

53. Willingham, S.B., Volkmer, J.P., Gentles, A.J., Sahoo, D., Dalerba, P., Mitra, S.S., Wang, J., Contreras-Trujillo, H., Martin, R., Cohen, J.D., et al. (2012). The CD47-signal regulatory protein alpha (SIRPa) interaction is a therapeutic target for human solid tumors. Proc Natl Acad Sci U S A 109, 6662–6667. 10.1073/pnas.1121623109.

54. Chen, C., Jing, W., Chen, Y., Wang, G., Abdalla, M., Gao, L., Han, M., Shi, C., Li, A., Sun, P., et al. (2022). Intracavity generation of glioma stem cell-specific CAR macrophages primes locoregional immunity for postoperative glioblastoma therapy. Sci Transl Med 14, eabn1128. 10.1126/scitranslmed.abn1128.

55. Jin, G., Chang, Y., and Bao, X. (2023). Generation of chimeric antigen receptor macrophages from human pluripotent stem cells to target glioblastoma. Immunooncol Technol 20, 100409. 10.1016/j.iotech.2023.100409.

56. Zhou, L., Song, Q., Zhang, X., Cao, M., Xue, D., Sun, Y., Mao, M., Li, X., Zhang, Z., Liu, J., and Shi, J. (2025). In vivo generation of CAR macrophages via the enucleated mesenchymal stem cell delivery system for glioblastoma therapy. Proc Natl Acad Sci U S A 122, e2426724122. 10.1073/pnas.2426724122.

57. Zhang, Y., Shen, J., Xu, Y., Feng, F., Han, X., Xi, K., Fang, Z., Zhang, Y., Wang, M., Wang, Z., et al. (2026). Engineered probiotics recruit CAR macrophages and establish immune memory to eradicate heterogeneous glioblastoma in mice. Cell Host Microbe 34, 212–229 e217. 10.1016/j.chom.2025.12.014.

58. Reiss, K.A., Angelos, M.G., Dees, E.C., Yuan, Y., Ueno, N.T., Pohlmann, P.R., Johnson, M.L., Chao, J., Shestova, O., Serody, J.S., et al. (2025). CAR-macrophage therapy for HER2-overexpressing advanced solid tumors: a phase 1 trial. Nat Med 31, 1171–1182. 10.1038/s41591-025-03495-z.

59. Li, X., Wang, X., Wang, H., Zuo, D., Xu, J., Feng, Y., Xue, D., Zhang, L., Lin, L., and Zhang, J. (2024). A clinical study of autologous chimeric antigen receptor macrophage targeting mesothelin shows safety in ovarian cancer therapy. J Hematol Oncol 17, 116. 10.1186/s13045-024-01635-5.

60. Wang, X., Wang, Y., Yu, L., Sakakura, K., Visus, C., Schwab, J.H., Ferrone, C.R., Favoino, E., Koya, Y., Campoli, M.R., et al. (2010). CSPG4 in cancer: multiple roles. Curr Mol Med 10, 419–429. 10.2174/156652410791316977.

61. Beard, R.E., Zheng, Z., Lagisetty, K.H., Burns, W.R., Tran, E., Hewitt, S.M., Abate-Daga, D., Rosati, S.F., Fine, H.A., Ferrone, S., et al. (2014). Multiple chimeric antigen receptors successfully target chondroitin sulfate proteoglycan 4 in several different cancer histologies and cancer stem cells. J Immunother Cancer 2, 25. 10.1186/2051-1426-2-25.

62. Hafner, C., Breiteneder, H., Ferrone, S., Thallinger, C., Wagner, S., Schmidt, W.M., Jasinska, J., Kundi, M., Wolff, K., Zielinski, C.C., et al. (2005). Suppression of human melanoma tumor growth in SCID mice by a human high molecular weight-melanoma associated antigen (HMW-MAA) specific monoclonal antibody. Int J Cancer 114, 426–432. 10.1002/ijc.20769.

63. Belote, R.L., Le, D., Maynard, A., Lang, U.E., Sinclair, A., Lohman, B.K., Planells-Palop, V., Baskin, L., Tward, A.D., Darmanis, S., and Judson-Torres, R.L. (2021). Human melanocyte development and melanoma dedifferentiation at single-cell resolution. Nat Cell Biol 23, 1035–1047. 10.1038/s41556-021-00740-8.

64. Jerby-Arnon, L., Shah, P., Cuoco, M.S., Rodman, C., Su, M.J., Melms, J.C., Leeson, R., Kanodia, A., Mei, S., Lin, J.R., et al. (2018). A Cancer Cell Program Promotes T Cell Exclusion and Resistance to Checkpoint Blockade. Cell 175, 984–997 e924. 10.1016/j.cell.2018.09.006.

65. Tirosh, I., Venteicher, A.S., Hebert, C., Escalante, L.E., Patel, A.P., Yizhak, K., Fisher, J.M., Rodman, C., Mount, C., Filbin, M.G., et al. (2016). Single-cell RNA-seq supports a developmental hierarchy in human oligodendroglioma. Nature 539, 309–313. 10.1038/nature20123.

66. Pellegatta, S., Savoldo, B., Di Ianni, N., Corbetta, C., Chen, Y., Patane, M., Sun, C., Pollo, B., Ferrone, S., DiMeco, F., et al. (2018). Constitutive and TNFalpha-inducible expression of chondroitin sulfate proteoglycan 4 in glioblastoma and neurospheres: Implications for CAR-T cell therapy. Sci Transl Med 10. 10.1126/scitranslmed.aao2731.

67. Al-Mayhani, M.T., Grenfell, R., Narita, M., Piccirillo, S., Kenney-Herbert, E., Fawcett, J.W., Collins, V.P., Ichimura, K., and Watts, C. (2011). NG2 expression in glioblastoma identifies an actively proliferating population with an aggressive molecular signature. Neuro Oncol 13, 830–845. 10.1093/neuonc/nor088.

68. Poli, A., Wang, J., Domingues, O., Planaguma, J., Yan, T., Rygh, C.B., Skaftnesmo, K.O., Thorsen, F., McCormack, E., Hentges, F., et al. (2013). Targeting glioblastoma with NK cells and mAb against NG2/CSPG4 prolongs animal survival. Oncotarget 4, 1527–1546. 10.18632/oncotarget.1291.

69. Wang, J., Svendsen, A., Kmiecik, J., Immervoll, H., Skaftnesmo, K.O., Planaguma, J., Reed, R.K., Bjerkvig, R., Miletic, H., Enger, P.O., et al. (2011). Targeting the NG2/CSPG4 proteoglycan retards tumour growth and angiogenesis in preclinical models of GBM and melanoma. PLoS One 6, e23062. 10.1371/journal.pone.0023062.

70. Greiner, D., Xue, Q., Waddell, T.Q., Kurudza, E., Chaudhary, P., Belote, R.L., Dotti, G., Judson-Torres, R.L., Reeves, M.Q., Cheshier, S.H., and Roh-Johnson, M. (2025). Human CSPG4-targeting CAR-macrophages inhibit melanoma growth. Oncogene. 10.1038/s41388-025-03332-0.

71. Kochenderfer, J.N., Feldman, S.A., Zhao, Y., Xu, H., Black, M.A., Morgan, R.A., Wilson, W.H., and Rosenberg, S.A. (2009). Construction and preclinical evaluation of an anti-CD19 chimeric antigen receptor. J Immunother 32, 689–702. 10.1097/CJI.0b013e3181ac6138.

72. Morrissey, M.A., Williamson, A.P., Steinbach, A.M., Roberts, E.W., Kern, N., Headley, M.B., and Vale, R.D. (2018). Chimeric antigen receptors that trigger phagocytosis. Elife 7. 10.7554/eLife.36688.

73. Geldres, C., Savoldo, B., Hoyos, V., Caruana, I., Zhang, M., Yvon, E., Del Vecchio, M., Creighton, C.J., Ittmann, M., Ferrone, S., and Dotti, G. (2014). T lymphocytes redirected against the chondroitin sulfate proteoglycan-4 control the growth of multiple solid tumors both in vitro and in vivo. Clin Cancer Res 20, 962–971. 10.1158/1078-0432.CCR-13-2218.

74. Rivera, Z., Ferrone, S., Wang, X., Jube, S., Yang, H., Pass, H.I., Kanodia, S., Gaudino, G., and Carbone, M. (2012). CSPG4 as a target of antibody-based immunotherapy for malignant mesothelioma. Clin Cancer Res 18, 5352–5363. 10.1158/1078-0432.CCR-12-0628.

75. Greiner, D., Scott, T.M., Olson, G.S., Aderem, A., Roh-Johnson, M., and Johnson, J.S. (2022). Genetic Modification of Primary Human Myeloid Cells to Study Cell Migration, Activation, and Organelle Dynamics. Curr Protoc 2, e514. 10.1002/cpz1.514.

76. Kidwell, C.U., Casalini, J.R., Pradeep, S., Scherer, S.D., Greiner, D., Bayik, D., Watson, D.C., Bae, D.H., Olson, G.S., Lathia, J.D., et al. (2023). Transferred mitochondria act as a signaling source, promoting proliferation. In U.o. Utah, ed.

77. Hsu, A.P., Donko, A., Arrington, M.E., Swamydas, M., Fink, D., Das, A., Escobedo, O., Bonagura, V., Szabolcs, P., Steinberg, H.N., et al. (2019). Dominant activating RAC2 mutation with lymphopenia, immunodeficiency, and cytoskeletal defects. Blood 133, 1977–1988. 10.1182/blood-2018-11-886028.

78. Mishra, A.K., Rodriguez, M., Torres, A.Y., Smith, M., Rodriguez, A., Bond, A., Morrissey, M.A., and Montell, D.J. (2023). Hyperactive Rac stimulates cannibalism of living target cells and enhances CAR-M-mediated cancer cell killing. Proc Natl Acad Sci U S A 120, e2310221120. 10.1073/pnas.2310221120.

79. Singh, M.M., Johnson, B., Venkatarayan, A., Flores, E.R., Zhang, J., Su, X., Barton, M., Lang, F., and Chandra, J. (2015). Preclinical activity of combined HDAC and KDM1A inhibition in glioblastoma. Neuro Oncol 17, 1463–1473. 10.1093/neuonc/nov041.

80. Gonzalez, M.A., Lu, D.R., Yousefi, M., Kroll, A., Lo, C.H., Briseno, C.G., Watson, J.E.V., Novitskiy, S., Arias, V., Zhou, H., et al. (2023). Phagocytosis increases an oxidative metabolic and immune suppressive signature in tumor macrophages. J Exp Med 220. 10.1084/jem.20221472.

81. Zhang, C.R., Xiong, L., Gu, M., Yeh, C.Y., Jerby, L., Kundaje, A., Greenleaf, W.J., and Bassik, M.C. (2025). Cancer cell phagocytosis induces an anti-inflammatory gene regulatory program in macrophages. bioRxiv. 10.1101/2025.08.18.670953.

82. Zhang, L., Tian, L., Dai, X., Yu, H., Wang, J., Lei, A., Zhu, M., Xu, J., Zhao, W., Zhu, Y., et al. (2020). Pluripotent stem cell-derived CAR-macrophage cells with antigen-dependent anti-cancer cell functions. J Hematol Oncol 13, 153. 10.1186/s13045-020-00983-2.

83. Klichinsky, M., Ruella, M., Shestova, O., Lu, X.M., Best, A., Zeeman, M., Schmierer, M., Gabrusiewicz, K., Anderson, N.R., Petty, N.E., et al. (2020). Human chimeric antigen receptor macrophages for cancer immunotherapy. Nat Biotechnol 38, 947–953. 10.1038/s41587-020-0462-y.

84. Glioblastoma Multiforme Treatment Market Analysis By Treatment (Chemotherapy, Immunotherapy, Targeted Therapy, Radiation Therapy, Other Treatments), By Drug Class (Temozolomide, Lomustine, Bevacizumab, Other Drug Classes), By End-use (Hospitals, Clinics, Ambulatory Surgical Centers), By Region and Companies - Industry Segment Outlook, Market Assessment, Competition Scenario, Trends and Forecast 2024-2033. (2024). https://market.us/report/glioblastoma-multiforme-treatment-market/.

85. Valon, L., Davidovic, A., Levillayer, F., Villars, A., Chouly, M., Cerqueira-Campos, F., and Levayer, R. (2021). Robustness of epithelial sealing is an emerging property of local ERK feedback driven by cell elimination. Dev Cell 56, 1700–1711 e1708. 10.1016/j.devcel.2021.05.006.

86. Toyama, Y., Peralta, X.G., Wells, A.R., Kiehart, D.P., and Edwards, G.S. (2008). Apoptotic force and tissue dynamics during Drosophila embryogenesis. Science 321, 1683–1686. 10.1126/science.1157052.

87. Gorelick-Ashkenazi, A., Weiss, R., Sapozhnikov, L., Florentin, A., Tarayrah-Ibraheim, L., Dweik, D., Yacobi-Sharon, K., and Arama, E. (2018). Caspases maintain tissue integrity by an apoptosis-independent inhibition of cell migration and invasion. Nat Commun 9, 2806. 10.1038/s41467-018-05204-6.

88. Elkington, P.T., Green, J.A., and Friedland, J.S. (2009). Analysis of matrix metalloproteinase secretion by macrophages. Methods Mol Biol 531, 253–265. 10.1007/978-1-59745-396-7_16.

89. Vinnakota, K., Zhang, Y., Selvanesan, B.C., Topi, G., Salim, T., Sand-Dejmek, J., Jonsson, G., and Sjolander, A. (2017). M2-like macrophages induce colon cancer cell invasion via matrix metalloproteinases. J Cell Physiol 232, 3468–3480. 10.1002/jcp.25808.

90. Wahl, S.M., McCartney-Francis, N., Allen, J.B., Dougherty, E.B., and Dougherty, S.F. (1990). Macrophage production of TGF-beta and regulation by TGF-beta. Ann N Y Acad Sci 593, 188–196. 10.1111/j.1749-6632.1990.tb16111.x.

91. Xing, X., Wang, Y., Zhang, X., Gao, X., Li, M., Wu, S., Zhao, Y., Chen, J., Gao, D., Chen, R., et al. (2021). Matrix stiffness-mediated effects on macrophages polarization and their LOXL2 expression. FEBS J 288, 3465–3477. 10.1111/febs.15566.

92. Friedman-DeLuca, M., Karagiannis, G.S., Condeelis, J.S., Oktay, M.H., and Entenberg, D. (2024). Macrophages in tumor cell migration and metastasis. Front Immunol 15, 1494462. 10.3389/fimmu.2024.1494462.

93. Liu, X., Lv, Z., Zou, J., Liu, X., Ma, J., Sun, C., Sa, N., and Xu, W. (2016). Elevated AEG- 1 expression in macrophages promotes hypopharyngeal cancer invasion through the STAT3-MMP-9 signaling pathway. Oncotarget 7, 77244–77256. 10.18632/oncotarget.12886.

94. Zhang, F., Wang, Z., Fan, Y., Xu, Q., Ji, W., Tian, R., and Niu, R. (2015). Elevated STAT3 Signaling-Mediated Upregulation of MMP-2/9 Confers Enhanced Invasion Ability in Multidrug-Resistant Breast Cancer Cells. Int J Mol Sci 16, 24772–24790. 10.3390/ijms161024772.

95. Yamanaka, N., Morisaki, T., Nakashima, H., Tasaki, A., Kubo, M., Kuga, H., Nakahara, C., Nakamura, K., Noshiro, H., Yao, T., et al. (2004). Interleukin 1beta enhances invasive ability of gastric carcinoma through nuclear factor-kappaB activation. Clin Cancer Res 10, 1853–1859. 10.1158/1078-0432.ccr-03-0300.

