## Supplemental Figures & Legends for "NG2-targeting macrophages inhibit 3D invasion of patient-derived glioblastoma spheroids"

### SUPPLEMENTAL FIGURES AND LEGENDS

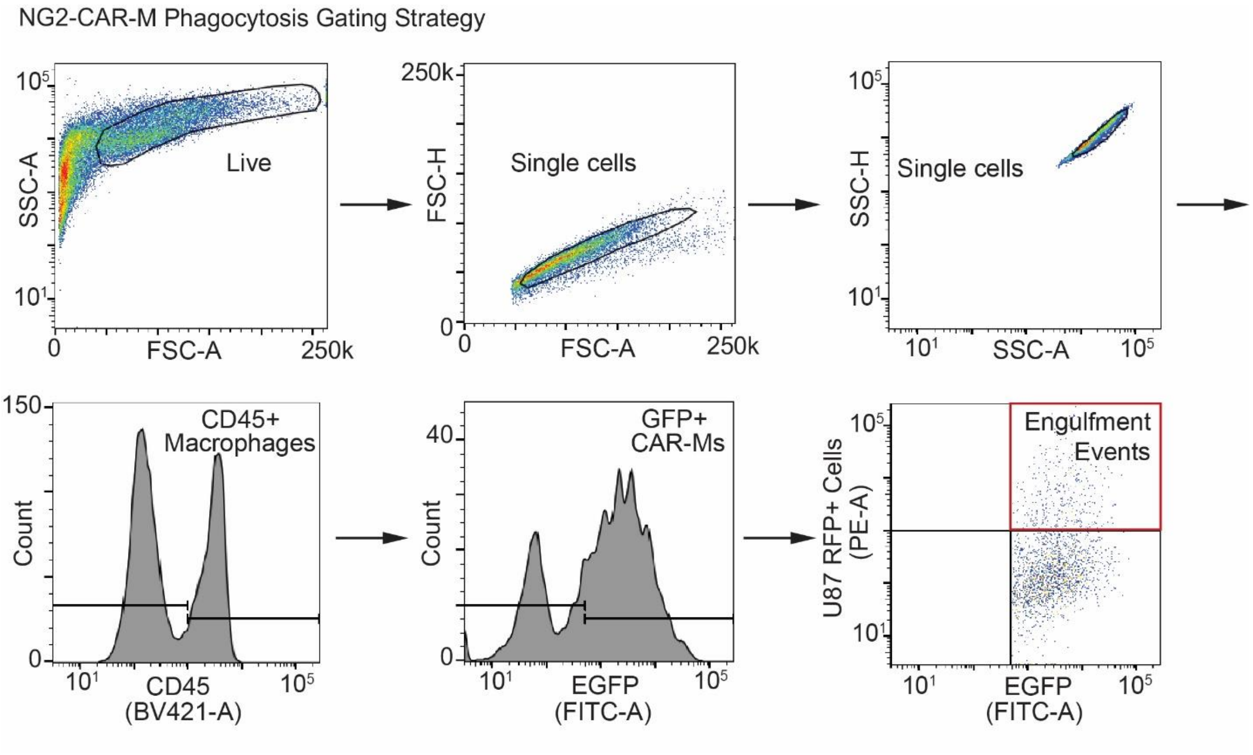


**Figure S1: NG2-CAR-Ms engulf GBM cells.** Representative gating strategy flow cytometry plots for NG2-CAR-Ms co-cultured with RFP+ GBM$^{U87}$ spheroids and gating for "Engulfment Events", as defined by CD45+ EGFP+ CAR-Ms positive for RFP+.

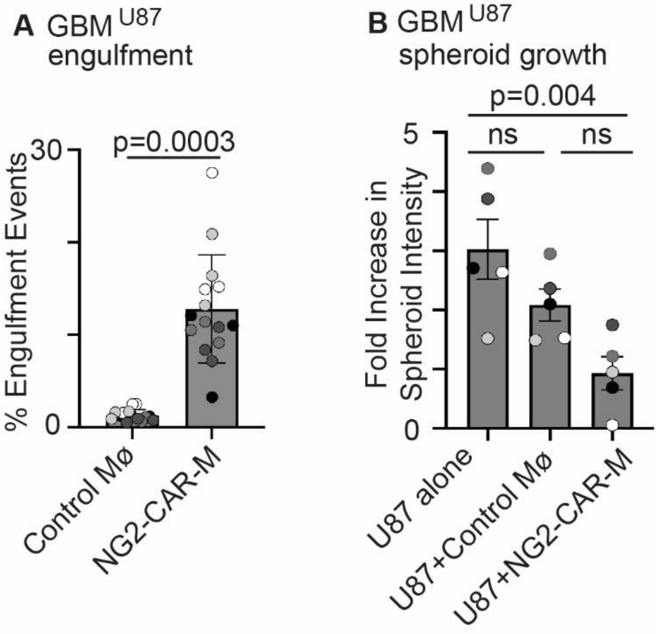


**Figure S2: NG2 CAR-M engulf and inhibit the growth of GBM$^{U87}$ cells**. A. Quantification of CAR-M-mediated engulfment events from Figure 2C and the raw data that corresponds to Figure 2D. n=4 independent PBMC donors (biological replicates) each as a shade of gray. Technical replicates are also shown for each biological replicate. mean +/- SEM, Two-tailed nested T-test. B. Raw data that corresponds to Figure 3C. Quantification of data in Figure 3B at 184 hours across conditions (n=5 independent PBMC donors (biological replicates), each with 6 technical replicates for every condition). Fluorescence intensity measurements are shown as a fold change at the last time point relative to the first time point. Biological replicates are shown as shades of gray, with technical replicates shown. mean +/- SEM, nested 1-way ANOVA with Tukey's multiple comparison test.

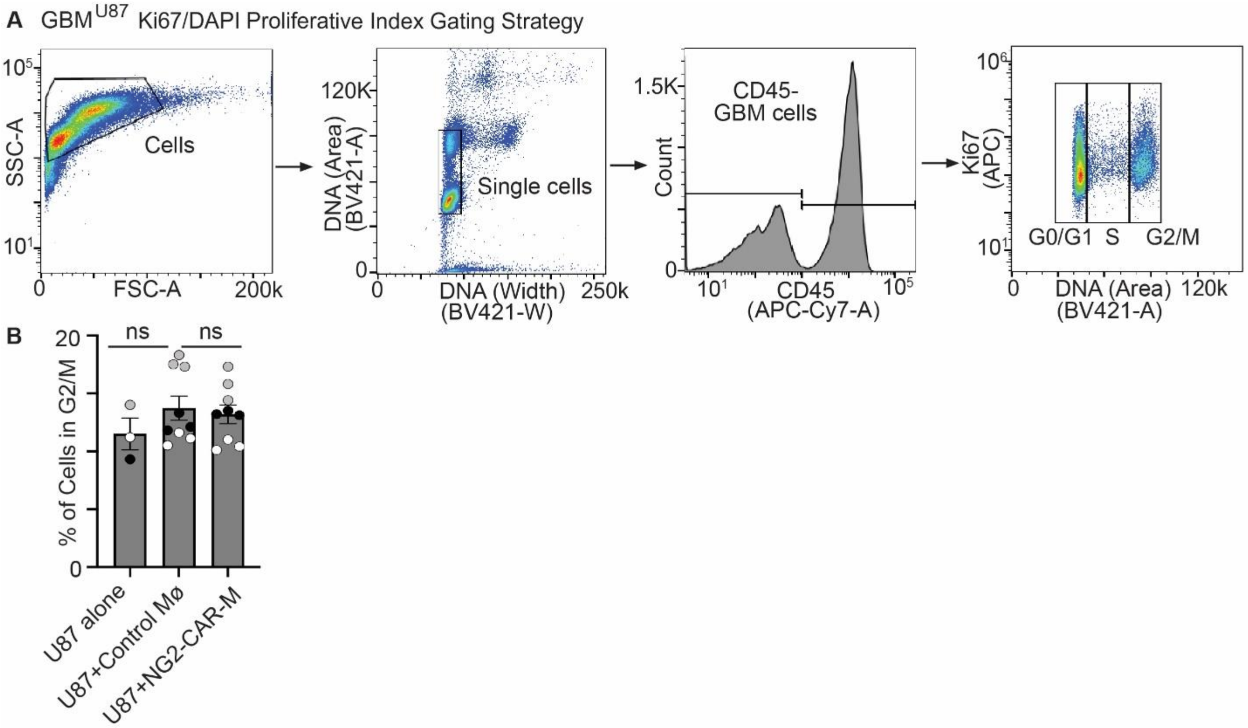


**Figure S3: GBM$^{U87}$ cells do not exhibit changes in proliferative capacity in the presence of NG2-CAR-Ms.** A. Representative gating strategy for flow cytometry plots for NG2-CAR-M/GBM$^{U87}$ spheroid co-cultures stained for Ki67 content to assign GBM$^{U87}$ cells into stages of the cell cycle. B. Raw data that corresponds to Figure 3E. Quantification of the percent of GBM$^{U87}$ cells in the G2/M phase of the cell cycle when cultured alone or with control macrophages or NG2-targeting CAR-Ms. n=3 independent PBMC donors each as a color (biological replicates), with each technical replicate shown. mean +/- SEM, nested 1-way ANOVA with Tukey's multiple comparison test.

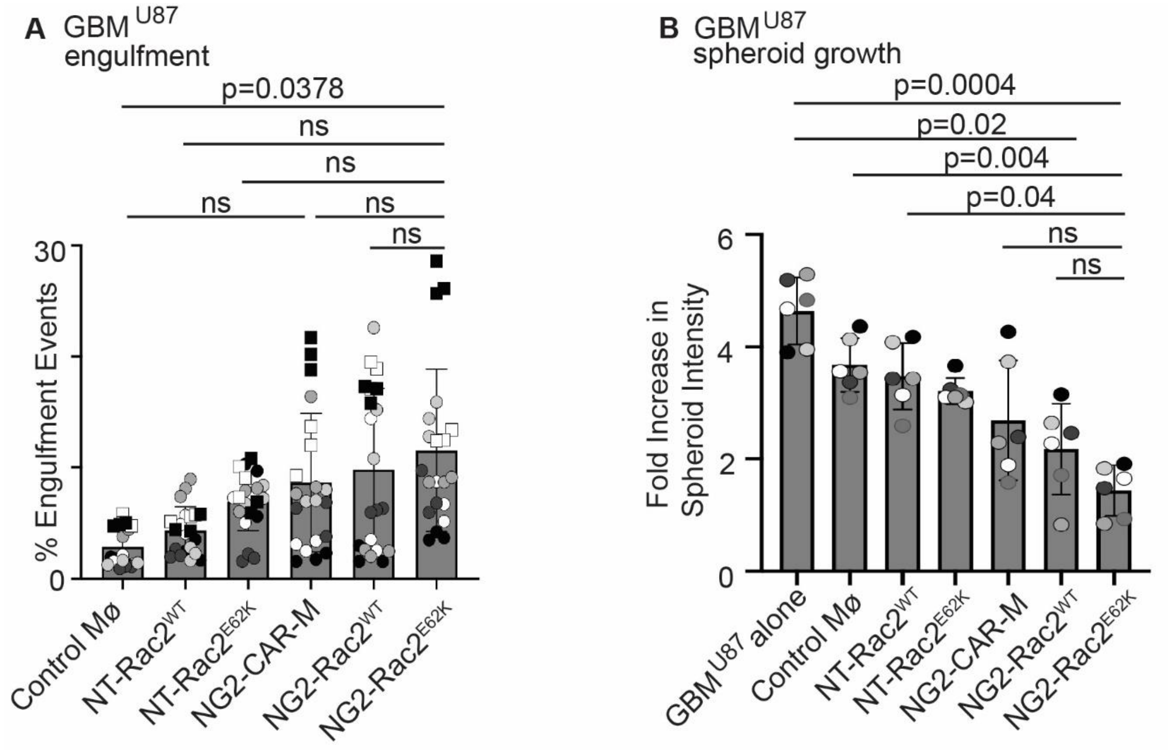


**Figure S4: NG2-Rac2$^{E62K}$ CAR-Ms exhibit higher rates of GBM$^{U87}$ engulfment and higher inhibition of spheroid growth.** A. Raw data that corresponds to Figure 4B. Quantification of EGFP+ CAR-M-mediated engulfment of RFP+ GBM$^{U87}$ cells by flow cytometry after 24 hours. n=7 independent PBMC donors (biological replicates). mean +/- SEM, nested 1-way ANOVA with Tukey's multiple comparison test. B. Raw data that corresponds to Figure 4D. Quantification of data in Figure 4C at day 10 across conditions (n=6 independent PBMC donors (biological replicates), each with 6 technical replicates for every condition). Data are shown as fold increase in spheroid fluorescence intensity for each biological replicate +/- SEM, nested 1-way ANOVA with Tukey's multiple comparison test.

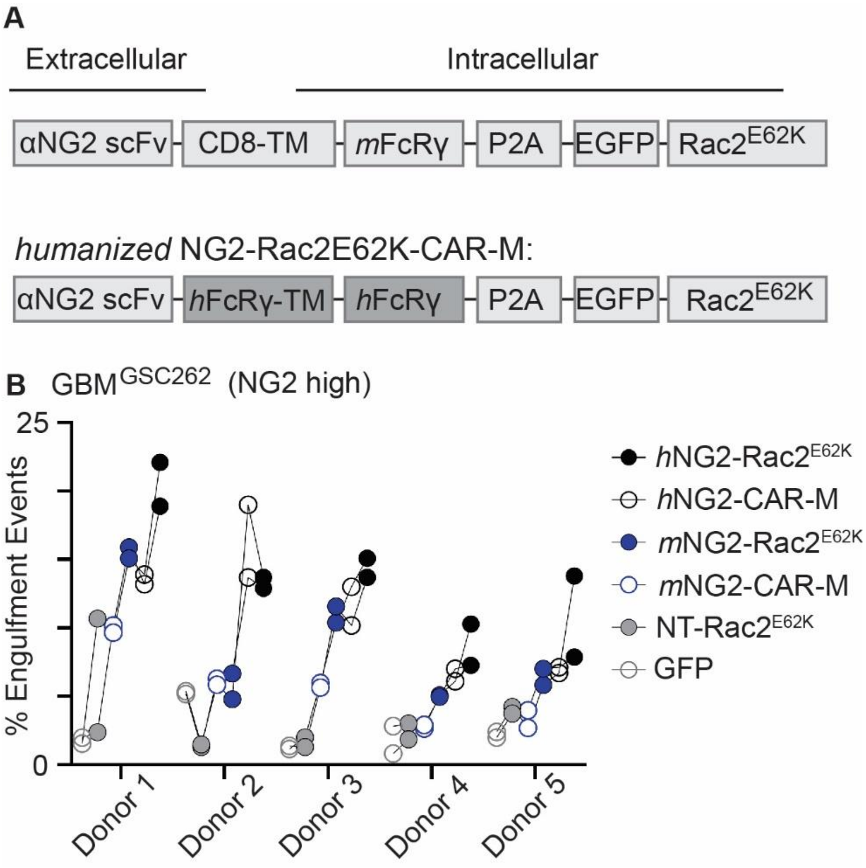


**Figure S5: Humanized NG2-CAR-Ms exhibit superior GBM$^{GSC262}$ engulfment compared to murine sequences.** A. Schematic of humanized NG2-CAR-M design (bottom, highlighted in darker gray and annotated with an "h") compared to murine sequences (top, annotated with an "m"). The transmembrane domain (TM) was also changed from the transmembrane sequence of human CD8 to the transmembrane sequence of human FcRgamma. B. Quantification of EGFP+ CAR-M-mediated engulfment of RFP+ GBM$^{GSC262}$ cells by flow cytometry after 24 hours, comparing murine CAR-Ms (annotated with an 'm") to humanized CAR-Ms (annotated with an "h"). Each line shows two technical replicates for each of the 5 PBMC donors, showing that hNG2-Rac2$^{E62K}$ CAR-Ms (black circles) showed higher engulfment capacity compared to all other CAR-Ms in 4 out of the 5 PBMC donors.

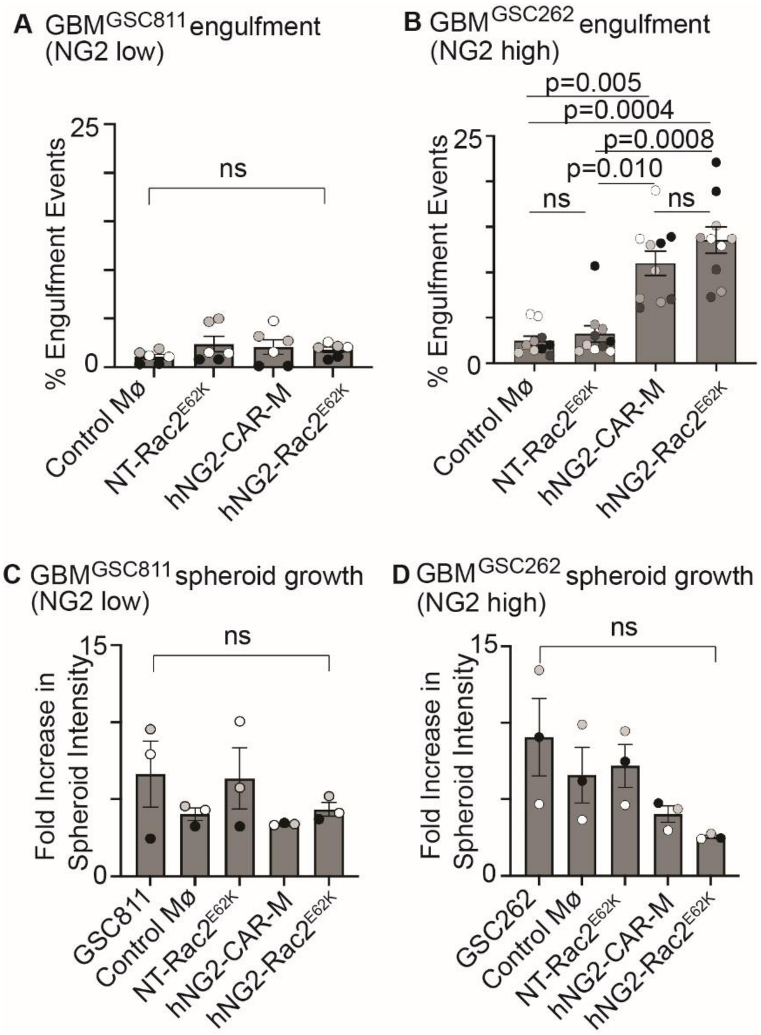


**Figure S6: NG2-CAR-Ms engulf patient-derived glioblastoma stem cells and inhibit their growth.** A. Raw data for Figure 5C. Quantification of EGFP+ CAR-M-mediated engulfment of mCherry+ GBM$^{GSC811}$ cells by flow cytometry after 24 hours. n=3 independent PBMC donors (biological replicates). mean +/- SEM, nested 1-way ANOVA with Tukey's multiple comparison test. B. Raw data for Figure 5D. Quantification (right) of EGFP+ CAR-M-mediated engulfment of mCherry+ GBM$^{GSC262}$ cells by flow cytometry after 24 hours. n=5 independent PBMC donors (biological replicates). mean +/- SEM, nested 1-way ANOVA with Tukey's multiple comparison test. C. Raw data for Figure 5E. Quantification of data at day 10 across conditions (n=6 independent PBMC donors (biological replicates), each with 6 technical replicates for every condition). Data are shown as fold increase in spheroid area. mean for each biological replicate +/- SEM, nested 1-way ANOVA with Tukey's multiple comparison test. D. Raw data for Figure 5F. Quantification of data at day 10 across conditions (n=3 independent PBMC donors (biological replicates), each with 6 technical replicates for every condition). Data are shown as fold increase in spheroid area. mean for each biological replicate +/- SEM, nested 1-way ANOVA with Tukey's multiple comparison test.

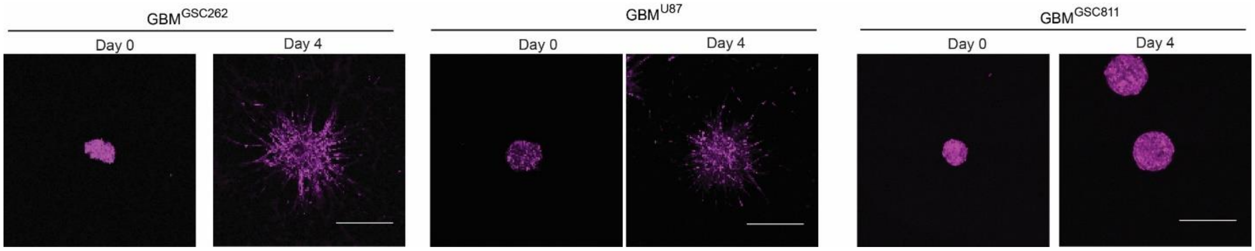


**Figure S7: GBM$^{GSC262}$ cells and GBM$^{U87}$ cells form spheroids that exhibit local invasion in 3D matrices**. Representative examples of spheroids formed from GBM$^{GSC262}$, GBM$^{U87}$, or GBM$^{GSC811}$ cells at day 0 and day 4. Scale bar = 500 $\mu$m.

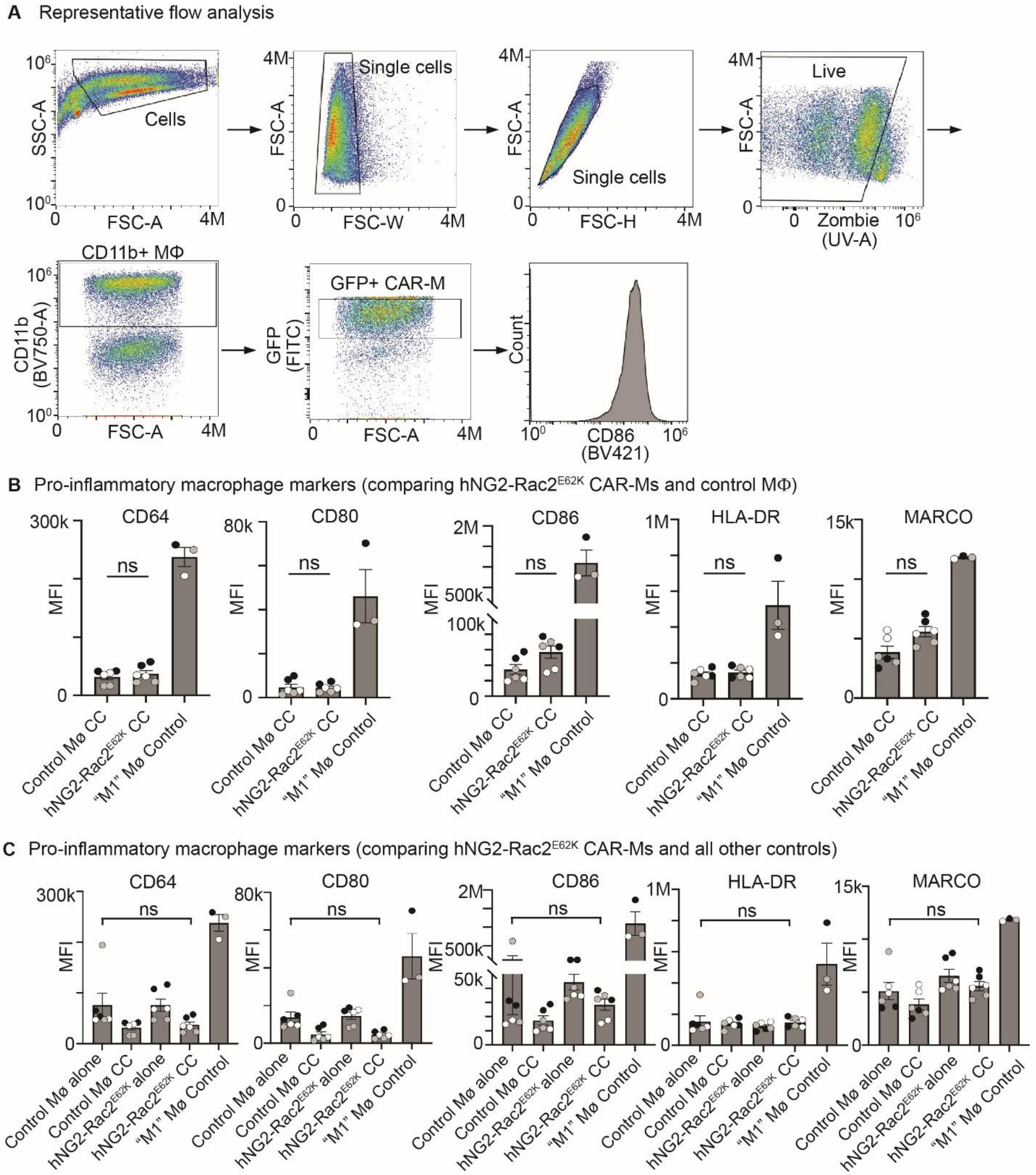


**Figure S8: NG2-Rac2$^{E62K}$ CAR-Ms do not exhibit differences in macrophage differentiation state compared to control macrophages.** A. Representative gating strategy for macrophage marker analysis. B. Quantification of median fluorescence intensity of pro-inflammatory macrophage markers (CD64, CD80, CD86, HLA-DR, and MARCO) comparing control macrophages versus hNG2-Rac2$^{E62K}$ CAR-Ms that were each in coculture ("CC") with GBM$^{GSC262}$ spheroids for 3 days ("Control MΦ CC" or "hNG2-Rac2$^{E62K}$ CC"). Macrophages stimulated with IFNγ/LPS for 3 days are used as a positive control for each marker ("M1 MΦ Control"). C. The same data in B are now graphed comparing macrophages cocultured with

GBM$^{GSC262}$ spheroids versus macrophages alone (no GBM cells) for control macrophages ("Control MΦ alone") and hNG2-Rac2$^{E62K}$ CAR-Ms ("hNG2-Rac2$^{E62K}$ alone"). For B,C: n=3 PBMC donors (biological replicates), each with 2 technical replicates (except the M1 MΦ Controls), with each biological replicate as a different color, nested t-test (for B) or nested 1-way ANOVA with Tukey's multiple comparison test (for C). M1 MΦ Controls were not included in the statistical tests as they were only used as a control for antibody staining.
